# A connectivity signature for glioblastoma

**DOI:** 10.1101/2021.11.07.465791

**Authors:** Ling Hai, Dirk C Hoffmann, Henriette Mandelbaum, Ruifan Xie, Jakob Ito, Erik Jung, Sophie Weil, Philipp Sievers, Varun Venkataramani, Daniel Dominguez Azorin, Kati Ernst, Denise Reibold, Rainer Will, Mario L. Suvà, Christel Herold-Mende, Felix Sahm, Frank Winkler, Matthias Schlesner, Wolfgang Wick, Tobias Kessler

**Author notes:** Corresponding author: Tobias Kessler MD, Neurology Clinic and Neurooncology Program at the National Center for Tumor Diseases & DKTK, DKFZ, Im Neuenheimer Feld 400, D-69120 Heidelberg, Germany. Equally contributing first authors (Ling Hai, Dirk C Hoffmann).

## Abstract

Tumor cell extensions called tumor microtubes (TMs) in glioma resemble neurites during neurodevelopment and connect glioma cells to a network that has considerable relevance for tumor progression and therapy resistance. The determination of interconnectivity in individual tumors has been challenging and the impact of tumor cell connectivity on patient survival remained unresolved so far. Here, a connectivity signature from single-cell RNA-sequenced (scRNA-Seq) xenografted primary glioblastoma (GB) cells was established and clinically validated. Thirty-four of 40 connectivity genes were related to neurogenesis, neural tube development or glioma progression, including the TM-network-relevant *GAP43* gene. Astrocytic-like and mesenchymal-like GB cells had the highest connectivity signature scores in scRNA-Seq data of patient-derived xenografts and patient samples. In 230 human GBs, high connectivity correlated with the mesenchymal expression subtype, *TP53* wildtype, and with dismal patient survival. *CHI3L1* was identified as a robust molecular marker of connectivity. Thus, the connectivity signature allows novel insights into brain tumor biology, provides a proof-of-principle that tumor cell connectivity is relevant for patients’ prognosis, and serves as a robust biomarker that can be used for future clinical trials.

**Statement of significance:** Integration of GB cells into functional networks drives tumor progression and resistance. Here, we established and validated a novel connectivity gene expression signature of single GB cells and whole tumors that can be easily applied to clinical and preclinical samples. It is shown that connectivity is determining prognosis combining molecular, functional and clinical insights into the disease.

## Introduction

Glioblastoma (GB) is the most common malignant primary brain tumor and patients have a median survival of about 15-20 months even when treated with full standard therapy (1). Resistance against new targeted approaches is pre-existing or acquired early and regularly, with no targeted therapy today that had proven efficacy in unselected studies (2). Tumor heterogeneity may play a major role in treatment resistance, as a subset of tumor cells might not be treatment sensitive, causing frequent and early relapses. Although not yet related to clinical resistance, different tumor cell populations have been detected by single-cell RNA sequencing (scRNA-Seq) techniques (3–5). Malignant cells in GB exist in at least four main cellular states that recapitulate distinct brain cell types, are influenced by the tumor microenvironment, and exhibit plasticity (5).

We have recently discovered that ultralong cellular protrusions named tumor microtubes (TMs) connect about half of the tumor cells to a multicellular network in patient samples of GB and preclinical models (6). Integration into these networks promotes resistance against radiotherapy (6), chemotherapy and surgical lesions (7). Until today, these TM networks have also been detected in incurable pediatric glioma types (8). TM networks facilitate long range communication of glioma cells by intercellular calcium waves, which is used for directed tumor self-repair, and a better cellular homeostasis (9, 10). TM networks receive synaptic neuronal input that activates glioma network communication, further driving glioma invasion and proliferation (8, 11). Tumor network connectivity appeared however variable between individual tumors (6) and the degree of connectivity relevant for the degree of resistance. Improved molecular understanding of connectivity to unravel candidate structures for intervention and the detection and quantification of the degree of connectivity in difficult to assess patient samples would be necessary to develop and evaluate disconnecting therapies (10). Few molecular drivers for TMs and their networks have been identified so far (6, 12), and the relation to single cell heterogeneity in a tumor is unknown.

Here, we develop and validate a gene expression signature of tumor network connectivity using a functional intravital dye transfer approach with subsequent bulk RNA sequencing (RNA-Seq) and scRNA-Seq. Next to fundamental insights into the molecular features of TM network connectivity, the resulting connectivity signature score proved to be a straightforward, reliable, and prognostic biomarker for this central cellular underpinning of glioma malignancy.

## Results

### Development of a connectivity signature for GB

We first aimed to explore the transcriptomic landscape of TM-connected GB cells. Three patient derived glioblastoma cell lines (PDGCLs), “S24”, “T269” and “P3XX”, were tagged by green fluorescent protein (GFP) and xenografted into mouse brains (**Figure 1A**). All three PDGCLs were confirmed to form TMs and TM-networks in the xenografted mouse models (**Figure 1B**), thus reflecting the TM connectivity regularly seen in diffuse astrocytomas and GBs of patients (**Figure 1C**). To label the TM-connected tumor cells, we utilized sulforhodamine (SR) 101 based staining method (**Figure 1A**). SR101 is a red fluorescent dye that is transported via cell-to-cell connections and has been shown to preferentially label highly connected glioma cells after local (6) and systemic (11, 12) application. After intravenous injection of SR101, highly connected tumor cells showed higher SR101 staining intensity compared to lowly connected tumor cells in these mouse models confirming the validity of the SR101 model used (**Figure 1D-E**). Tumors were then harvested and subjected to fluorescence-activated cell sorting (FACS). The FACS-sorted SR101^high^ cells (highly connected tumor cells) and SR101^low^ cells (lowly connected tumor cells) were sequenced by RNA-Seq and scRNA-Seq (**Figure 1A, Supplementary Table 1**).

**Figure 1.**
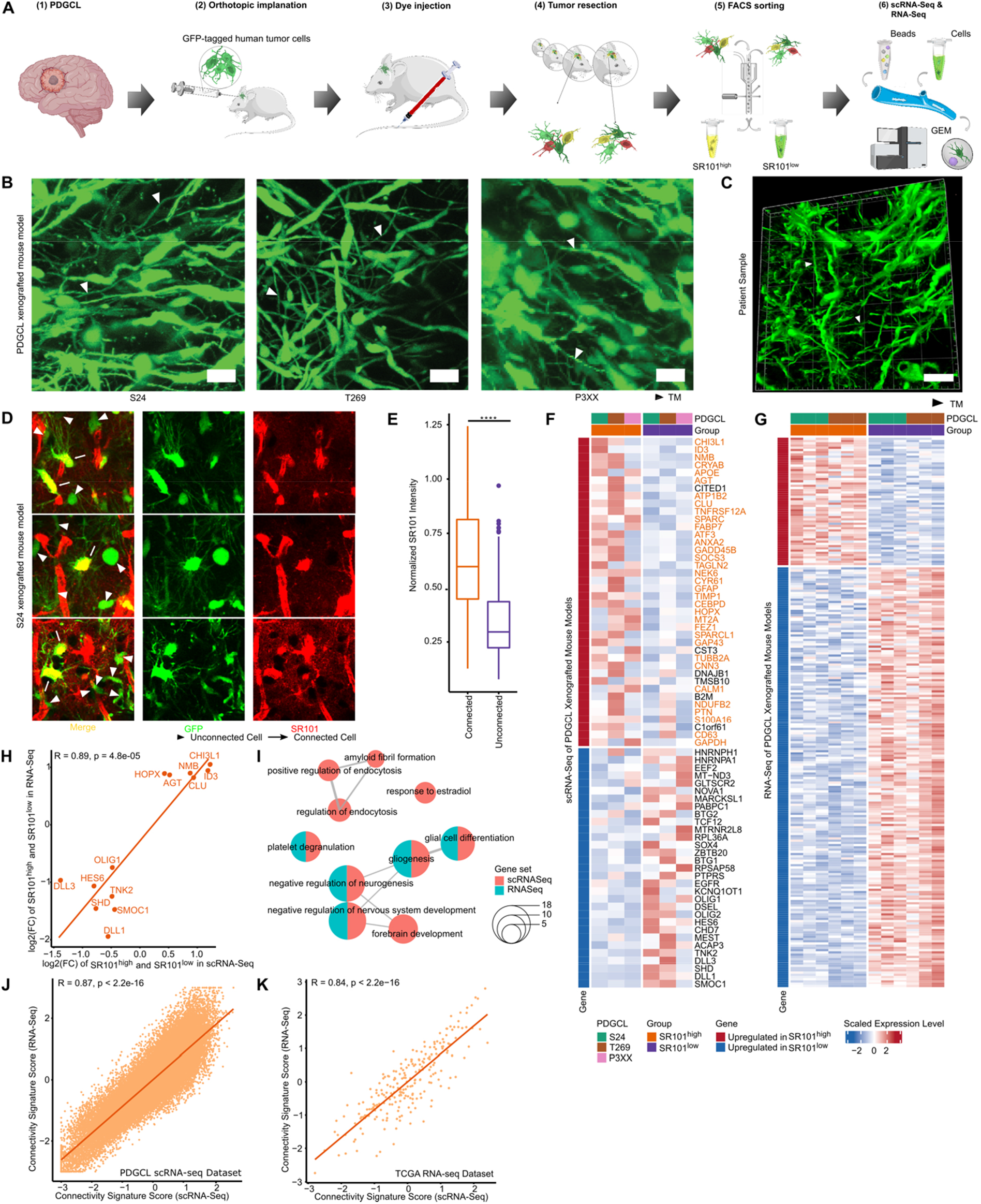
Development of the connectivity signature. **A,** Experimental design of the connectivity signature development. **B,** Representative two-photon microscopy images of three xenografted PDGCLs used for scRNA-seq. Arrowhead showing TMs. Scale bars, 20 µm. **C**, Representative confocal microscopy image from a GB patient in three dimensions. Arrowhead showing TMs. Scale bar, 20 µm. **D,** Representative two- photon microscopy images of S24 xenografted PDGCL; Red, SR101; Green, TurboGFP. Arrow mark showing highly connected cells and arrowhead showing lowly connected cells. **E,** Box plot of normalized SR101 intensity in 287 highly connected and 228 lowly connected cells from S24 xenografted PDGCL. P value was calculated by Mann-Whitney U test. ****, p < 0.0001. **F,** Heat map showing average expression levels of scRNA-Seq-derived 71 connectivity genes in SR101^high^ and SR101^low^ tumor cells from three xenografted PDGCLs. **G,** Heat map showing expression levels of RNA- Seq-derived 245 connectivity genes in SR101^high^ and SR101^low^ samples (n = 3) from two xenografted PDGCLs. Expression levels were normalized and batch effect removed. **H,** Scatter plot showing the log2 fold changes of SR101^high^ and SR101^low^ samples in scRNA-Seq and RNA-Seq datasets. 13 common connectivity genes were plotted. **I**, Enrichment map showing 10 most enriched GO biological processes in scRNA-Seq-derived gene set and/or RNA-Seq-derived gene set. The Pie charts with two colors indicating the consensus GOs between scRNA-Seq-derived and RNA-Seq- derived gene sets. The size of pie chart showing the number of overlapped genes between gene sets and GOs. **J-K,** Scatter plot showing connectivity signature scores based on connectivity genes derived from scRNA-Seq and RNA-Seq datasets. **J**, 35,822 cells from three xenografted PDGCLs scRNA-seq dataset. **K**, 230 samples from TCGA *IDH* wt GB RNA-Seq datasets. **F-G, J-K,** Values were scaled and centered across samples/cells, and winsorized to −3 and 3. **H, J-K,** Pearson correlation test was used to calculate correlation coefficients and p values. PDGCL, patient derived glioblastoma cell line.

To identify differentially expressed genes (DEGs) between highly and lowly connected tumor cells, multiple differential expression analyses (see **Methods**) were performed in the scRNA-Seq dataset of PDGCL xenografted mouse models. We obtained 71 DEGs that conserved in at least two PDGCL xenografted mouse models (**Figure 1F**). Among the 71 DEGs, 40 DEGs were found to be upregulated and 31 DEGs downregulated in highly connected tumor cells. 34 of 40 (85%) upregulated DEGs were previously described in the context of general cellular connectivity, but mainly not known in GB, development of neuronal cells, or as characteristic markers of GB progression (**Supplementary Table 2**). Of these, growth associated protein 43 (*GAP43)* has been characterized as a key player in TMs formation and TM-dependent cell-to-cell connectivity in gliomas (6) and apolipoprotein E (*APOE*) has recently been identified as a singular cluster marker for highly connected tumor cells (11).

For comparison to scRNA-Seq dataset, SR101^high^ and SR101^low^ tumor cells from two PDGCLs (“S24” and “T269”) were subjected to RNA-Seq. 245 DEGs were identified in RNA-Seq data (**Figure 1G**), of which, 13 DEGs were also identified in scRNA-Seq analyses with a high fold change correlation (R = 0.89, *p* = 4.8×10^-5^, **Figure 1H).**

Chitinase-3-like protein 1 (*CHI3L1*) was among the overlapped genes that showed the highest fold-change in both scRNA-Seq and RNA-Seq analyses **(Figure 1H)**. Remarkably, we have recently identified CHI3L1 as a key cerebrospinal fluid proteomic biomarker in GB in an independent study (13). The mRNA level and protein level of *CHI3L1* in GB were highly correlated (*n* = 93, R = 0.85, **Supplementary Figure 1A**). Previous analyses found an association of *CHI3L1* with the mesenchymal subtype in GB (14). Treatment of PDGCLs with CHI3L1 recombinant protein increased cell-to-cell connectivity and addition of a monoclonal antibody against CHI3L1 decreased cell-to-cell connectivity, arguing for a functional role of CHI3L1 in tumor cell connectivity (**Supplementary Figure 1B-C**). Furthermore, CHI3L1 was described to be involved in several cancer promoting mechanisms (15), therefore being one particularly interesting marker of highly connected tumor cells.

To compare the scRNA-Seq derived 71 DEGs and the RNA-Seq derived 245 DEGs on a gene set level, gene ontology (GO) enrichment analysis was performed. “Negative regulation of neurogenesis” and “Negative regulation of nervous system development” were part of the main enriched GO terms commonly in both gene sets (**Figure 1I**), which further supports the finding the TM-network formation follows neurodevelopmental mechanisms. The GO semantic similarity between these two gene sets were high (similarity = 0.814). In addition, we performed enrichment analysis against all gene pre-ranked by folder changes between SR101^high^ and SR101^low^ samples, and found that“Neurogenesis” was significantly upregulated in SR101^low^ samples in both scRNA-Seq and RNA-Seq datasets (**Supplementary Figure 2**).

Furthermore, we calculated scores (see **Methods**) based on the average expression levels of the RNA-Seq-derived or scRNA-Seq-derived gene set. The performances of both scores were tested against the labels of the SR101 FACS sorting. The score based on scRNA-Seq-derived gene set yielded incrementally higher accuracy compared to score based on RNA-Seq-derived gene set (0.83 vs. 0.79, **Supplementary Table 3**). However, the overall concordances between both scores tested on scRNA-Seq and RNA-Seq data were high (R = 0.87 and R = 0.84 respectively, **Figure 1J-K**). In addition, scores based on random generated gene sets used as negative controls resulted in expected poor performance (average accuracy = 0.49). Therefore, we decided to use the scRNA-Seq derived gene set as a connectivity signature for further evaluation. Hereafter, the term “*connectivity signature*” refers to the gene set of 71 connectivity related DEGs derived from scRNA-Seq of PDGCL xenografted mouse models, whereas the term “*connectivity signature score*” refers to a number calculated to describe the extent of connectivity based on the expression levels of the 71 DEGs.

### Two distinct GB cell subpopulations are characterized by high connectivity signature scores

In scRNA-Seq of three PDGCL xenografted mouse models, we obtained 35,822 tumor cells with a median of 5,686 cells per sample and 2,086 genes per cell. We confirmed higher connectivity signature scores in SR101^high^ compared to SR101^low^ tumor cells (**Figure 2A-B, Supplementary Figure 3A-C**). The connectivity signature genes *GAP43, APOE*, and *CHI3L1* had higher expression in the highly connected group (**Figure 2C**). Recent large single-cell studies have identified gene expression signatures that allow to identify distinct glioma cell states: astrocytic-like (AC), mesenchymal-like (MES), oligodendrocyte progenitor-like (OPC), and neuronal progenitor-like (NPC) (5). We applied these signatures on the scRNA-Seq data of PDGCL xenografted mouse models to associate connectivity signature scores with certain cell states (**Figure 2D**). Highly connected SR101^high^ tumor cells were predominantly assigned to the AC and MES cell states while lowly connected SR101^low^ tumor cells were mainly assigned to the NPC and OPC cell states (**Figure 2D-E**). The connectivity signature scores were higher in AC and MES1 cell states compared to NPC and OPC cell states (**Figure 2F**). An unexpectedly high degree of overlap was found between the connectivity signature genes and cell-state-defining genes, in particular in the AC and MES1 cell states (AC 10/40, 25%; MES1 7/51, 14%; MES2 2/51, 4%; NPC1 1/51, 2%, **Figure 2G**). Several cell-state-defining genes of AC and MES1 cell states (5) are tumor cell connectivity associated genes in GB, like connexin 43, also known as gap junction protein alpha 1 (*GJA1,* (6), tweety-homologue 1 (*TTYH1,* (16) and the correlative marker *APOE* (11). Of the 40 upregulated DEGs in highly connected tumor cells, a subset was primarily expressed in the AC or/and MES cells, while the 31 downregulated DEGs were expressed in OPC or/and NPC cells (**Figure 2H**). In summary, the SR101 methodology allowed us to not only provide a broad map of the transcriptomic properties of highly connected *versus* lowly connected GB cells, but also to link functional and molecular connectivity features to known distinct tumor cell subpopulations in GB.

**Figure 2.**
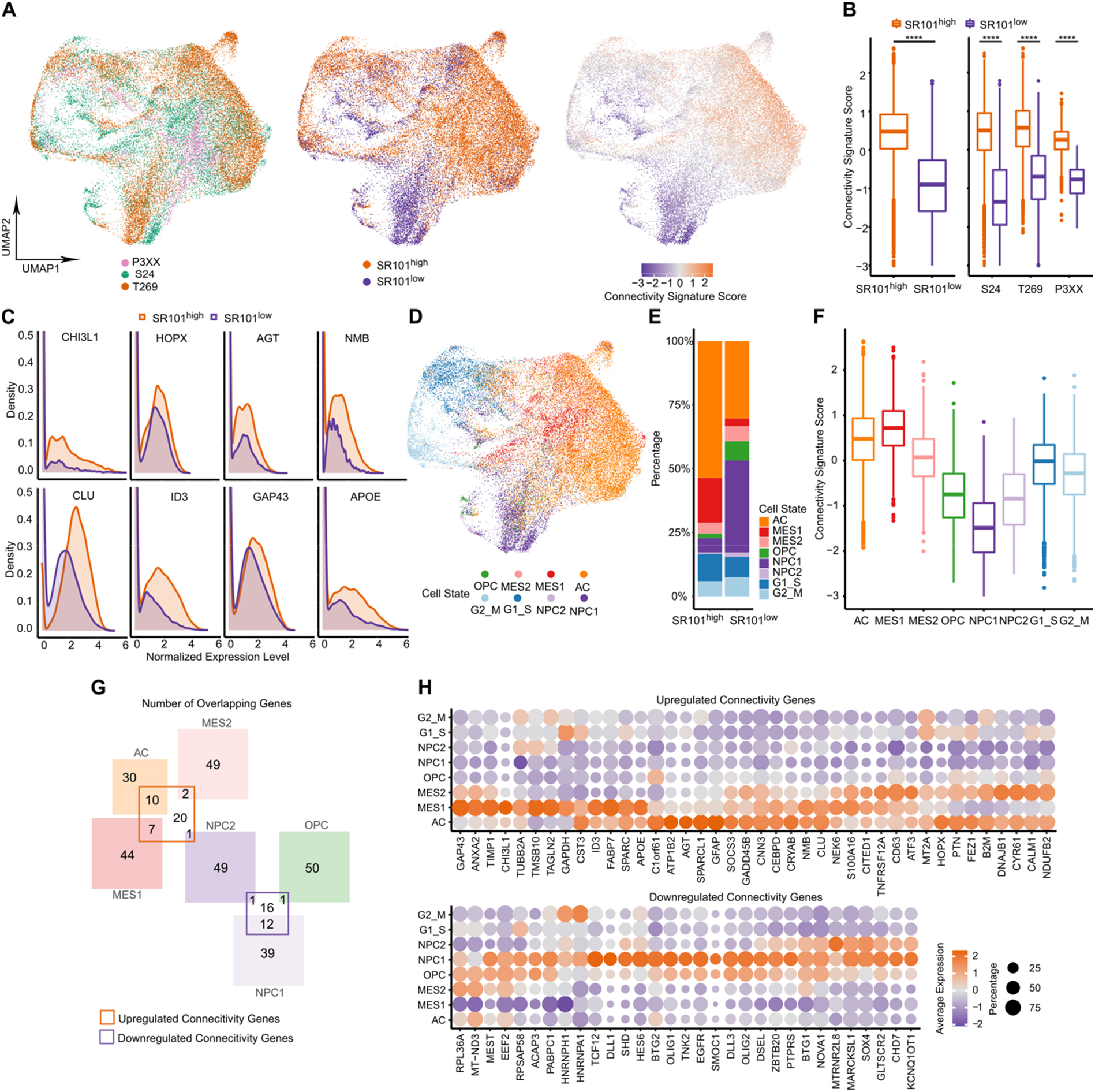
Connectivity signature scores in scRNA-seq of PDGCL xenografted mouse models. **A,** UMAPs of 35,822 cells in three xenografted PDGCLs scRNA-seq datasets. Left, colored by the derived PDGCL. Middle, colored by SR101-based sorting. Right, colored by connectivity signature scores. **B,** Box plot of connectivity signature scores in SR101^high^ and SR101^low^ cells. Left, cells from all three PDGCLs. Right, separated in each PDGCLs. P values were calculated by Mann-Whitney U test. ****, p < 0.0001.**C,** Density plot of normalized expression levels of genes in SR101^high^ and SR101^low^ cells. Upregulated common genes in scRNA-Seq-derived and RNA-Seq-derived connectivity genes (i.e., *CHI3L1, HOPX, AGT, NMB, CLU, ID3*) and the two upregulated scRNA-Seq-derived connectivity genes *APOE* and *GAP43* are shown. **D,** UMAP of cells in PDGCLs colored by cell states. **E,** Distribution of cell states in SR101^high^ and SR101^low^ cells. **F,** Box plot of connectivity signature scores in each cell state. P values in MES1 and MES2 were calculated by Mann-Whitney U test. **G,** Venn diagram showing the number of overlap genes between 71 connectivity genes and cell-state-defining genes. **H,** Dot plot of average expression levels of each connectivity gene in each cell state. Dot size indicates the frequency of cells that express the respective gene. Top, 40 upregulated connectivity genes in SR101^high^ cells. Bottom, 31 downregulated connectivity genes in SR101^high^ cells. **A, B, F,** Connectivity signature scores were scaled and centered across cells, and winsorized to −3 and 3.

### The connectivity signature score reflects true cell-to-cell connections in GB

To cross-validate that the connectivity signature score reflects actual morphological and physiological tumor cell connectivity, we first assessed its performance in induced connectivity of tumor cells experimentally *in vitro*. Four PDGCLs were subjected to different culture conditions: in the “TM+” condition, the cells adhered to the bottom of the flask and increasingly extended TMs, interconnecting single GB cells to tumor cell networks, while in the “TM-” condition, the cells formed floating spheroids and cells interconnections by TM were much less observed (**Figure 3A-B**).

**Figure 3.**
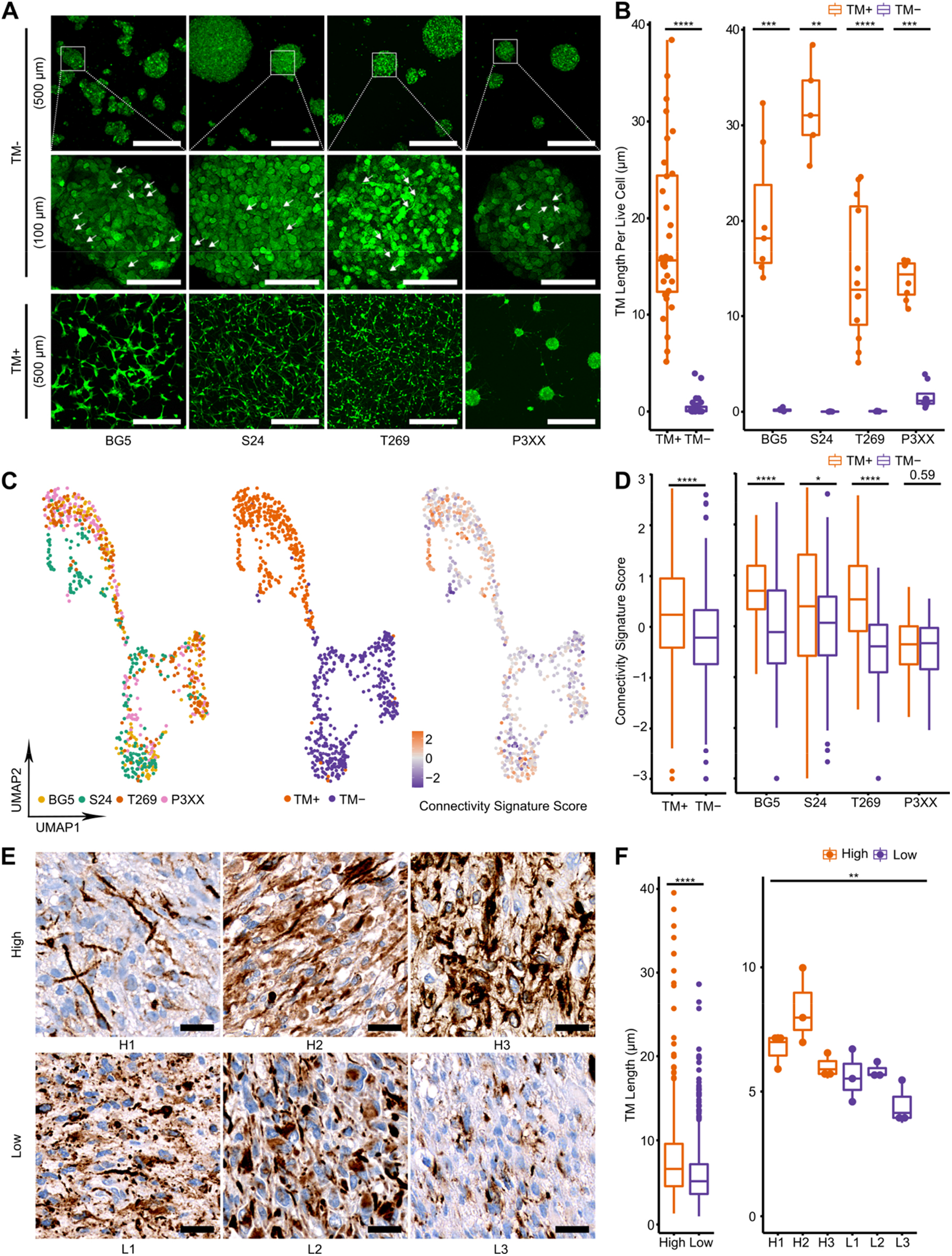
Connectivity signature scores reflected the true cell-to-cell connections. **A,** Representative confocal microscopy images from four PDGCLs. Top, cells cultured in TM- condition. Scale bars, 500 µm. Middle, zoom in from the top panel. Scale bars, 100 µm. Arrows indicate TMs. Bottom, cells cultured in TM+ condition. Scale bars, 500 µm. **B,** Box plot of the sum of TM lengths (µm) per live cell in 63 image crops of PDGCLs. Left, cells from all four PDGCLs. Right, separated in each PDGCL. **C,** UMAPs of 735 cells of the four PDGCLs. Left, colored by the derived PDGCL. Middle, colored by culturing conditions. Right, colored by connectivity signature scores. Scores were scaled and centered across cells, and winsorized to −3 and 3. **D,** Box plot of connectivity signature scores in cells. Left, cells from all four PDGCLs. Right, separated in each PDGCL. **E,** Representative immunohistochemistry staining images of TMs in six GB patients. Top, three patients with high connectivity signature scores (H1, H2 and H3). Bottom, three patients with low connectivity signature scores (L1, L2 and L3). Scale bars, 20 µm. **F,** Box plot of 898 TM lengths (µm) in patients. Left, TM lengths in all patients. Right, median of TM lengths per image crop in each patient (n = 3). **B, D, F,** P values in TM+ and TM- groups or high and low connectivity signature score groups were calculated by Mann-Whitney U test. *, p < 0.05; **, p < 0.01; ***, p < 0.001; ****, p < 0.0001.

We then performed scRNA-Seq of tumor cells from four PDGCLs cultured under TM+ and TM- conditions and obtained 735 cells with a median of 90 cells per sample and 4,893 genes per cell (**Supplementary Table 1**). PDGCLs cultured under the two conditions clustered separately from each other in a Uniform Manifold Approximation and Projection (UMAP) analysis (**Figure 3C**), with a higher connectivity signature score in TM+ cultured cells when compared to those cultured under TM- conditions (**Figure 3D**) confirming that experimental induction of TMs and their multicellular networks accompanied with an increase in the connectivity signature score. Furthermore, increased expression of three main markers of connectivity, *GAP43*, *APOE* and *CHI3L1* in TM+ cells was confirmed by qPCR (**Supplementary Figure 4A**).

Next, we aimed to validate the impact of the connectivity signature score in GB patient tumor tissues. We used immunofluorescence staining of nestin to assess connectivity in three patient tumor samples with high connectivity signature scores and three samples with low connectivity signature scores (**Figure 3E and Supplementary Figure 4B**). Samples with high connectivity signature scores showed increased length of TM-like structures compared to those with low connectivity signature scores (**Figure 3F**), which is a good estimation for TM connectivity in thin paraffin sections (6).

Together, this data supports the validity of the connectivity gene expression signature, both by *in vitro* assays and patient samples, confirming a meaningful interrelation of cellular and connectivity signature score-determined molecular connectivity.

### Applying the connectivity signature to GB patient samples

To test the performance of the connectivity signature in patient GB cells, 21 GB tumor samples were subjected to scRNA-Seq. All tumors were diagnosed as GB, isocitrate dehydrogenase *(IDH)* wildtype (wt), world health organization (WHO) grade 4, and the diagnoses were confirmed with methylation array analysis (**Supplementary Table 4**). A median of 11,192 cells per sample and 995 genes per cell passed quality control totaling in 213,444 single cells (**Figure 4A, Supplementary Figure 5A, Supplementary Table 4**).

**Figure 4.**
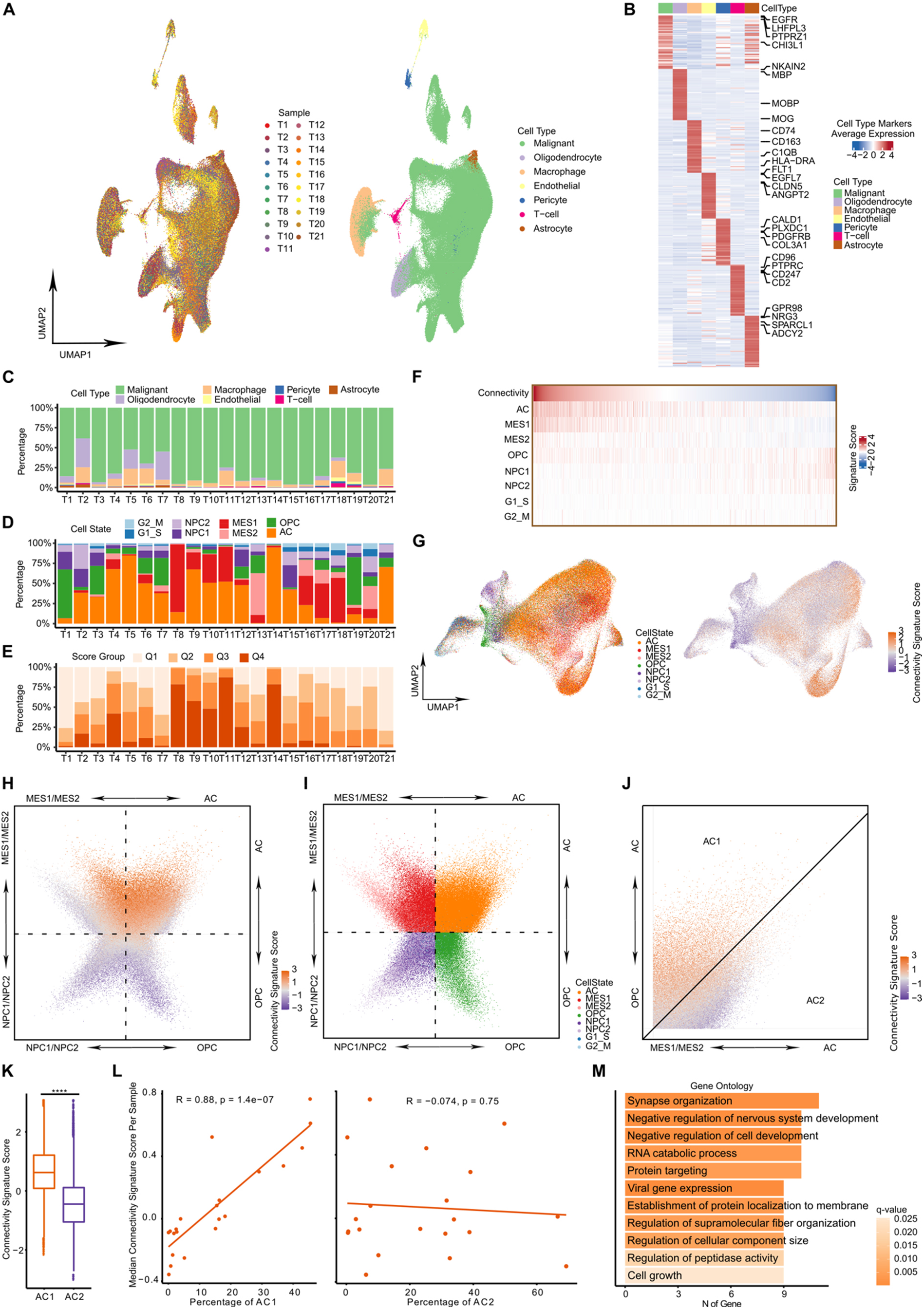
Connectivity signature scores in snRNA-Seq of patient samples. **A,** UMAP of 213,444 single cells from 21 GB patient samples. Left, colored by samples. Right, colored by cell types. **B,** Heat map of average expression levels of top 50 markers per cell type. Expression levels were scaled and centered across cell types, and winsorized to −3 and 3. **C,** Frequency of malignant and non-malignant cell types in each sample. **D,** Frequency of malignant cell states in each sample. **E,** Frequency of connectivity signature score group in each sample. Connectivity signature scores were grouped by four quartiles. **F,** Heat map showing connectivity signature scores and cell state signature scores in patient malignant cells. Each column represents one cell. **G,** UMAPs of patient malignant cells. Left, colored by cell states. Right, colored by connectivity signature scores. **H-J,** Two-dimensional representation of patient malignant cells according to cell state signature scores. **H,** colored by connectivity signature scores. **I,** colored by cell states. **J,** represented only AC cells (zoomed in from **H**). AC cells are separated by a line with slope 1 into AC1 and AC2 subtypes. **K,** Box plot of connectivity signature scores in AC1 and AC2 subtypes. P values in two groups were calculated by Mann-Whitney U test. ****, p < 0.0001. **L,** Scatter plot showing frequency of AC subtypes (Left, AC1; Right, AC2) and the median connectivity signature scores per patient sample. Dot indicates patient sample. 21 patient samples were shown. Pearson correlation test was used to calculate correlation coefficients and p values. **M,** The enriched GOs of 100 DEGs between AC1 and AC2 subtypes. Top 10 GOs were ordered by the number of genes overlapped between GO genes and DEGs. **F, G, H, J, K,** Signature scores were scaled and centered across cells, and winsorized to −3 and 3.

We classified malignant and non-malignant cells using previously defined marker genes (5,17,18) and copy number variation (CNV) analysis (**Figure 4A-C, Supplementary Figure 5A-D**). Within the malignant cells, the AC cells were predominant in most tumors although a high degree of heterogeneity in the cell states was observed between the tumors (**Figure 4D**). The connectivity signature score was also highly heterogeneous between tumors (**Figure 4E**), but consistently highest in AC and MES1 tumor cells and lowest in OPC and NPC like cells, confirming the results from the xenografted mouse models (**Figure 4F-I**).

Furthermore, the higher number of analyzed tumor cells allowed us to detect that AC tumor cells, similarly to the MES tumor cells, display two subgroups with higher (named AC1) and lower (named AC2) connectivity signature score (**Figure 4J-K**). The frequency of AC1 tumor cells in patient samples, but not the frequency of AC2, highly correlated with the median connectivity signature scores in each sample (**Figure 4L**). We analyzed DEGs between AC1 and AC2 cells and found *APOE* and *CHI3L1* to be significantly upregulated in AC1. GO term enrichment analysis on DEGs identified“synapse organization” as the GO term involving the largest number of DEGs (**Figure 4M**). This is in line with the recent discovery that glutamatergic neuron-glioma synapses do exist, mainly located on TMs, with the glioma cell as the principle postsynaptic partner, and with neuronal synaptic input which is strongly associated with glioma network activation (8, 11).

Here, we could validate in human GB samples that in particular tumor cells from two distinct subpopulations - AC1 and MES1 – are responsible for the establishment of cell-to-cell connectivity.

### The connectivity signature gene *CHI3L1* is a robust marker for connectivity

The analyses outlined previously suggested a role for *CHI3L1* in our connectivity signature. Therefore, we investigated the expression pattern of *CHI3L1* more deeply. *CHI3L1* is a conserved marker for SR101^high^ cells across all cell states. *CHI3L1* expression was highly correlated with connectivity signature scores in both The Cancer Genome Atlas (TCGA, *n* = 230) and Chinese Glioma Genome Atlas (CGGA, *n* = 141) *IDH* wt GB datasets (*r* = 0.73, *p* < 2.2*10^-16^, **Figure 5A**), which is the highest correlation of any single gene. Furthermore, higher *CHI3L1* expression was associated with worse overall survival in both patient datasets (**Figure 5B-C**). This effect was retained in a multivariate analysis adjusting for ages and genders (**Figure 5C**). High *CHI3L1* expression was found to be highly specific for GB compared to 30 other tumor types and related normal tissues (**Figure 5D**). Consistently, in our scRNA-Seq data of patient samples, *CHI3L1* expression was high in the high connectivity AC1 and MES1 tumor cell populations, but low in low connectivity tumor cell populations as well as non-malignant astrocytes, oligodendrocytes, vascular and immune cells (**Figure 5E**). To test whether CHI3L1 expressed areas are directly associated with cell-to-cell connected areas, we used tumors with high connectivity signature scores and long protrusions, and tumors with low connectivity signature scores and short protrusions. In tumors with long protrusions and high connectivity signature scores, we measured higher CHI3L1 protein levels assessed by immunohistochemistry for each sample (**Figure 5F-H**). In particular, even in heterogenous tumors, the correlation of CHI3L1 staining intensity and TM length assessment was also valid within the matched crops in adjacent sections (**Supplementary Figures 6-7**).

**Figure 5.**
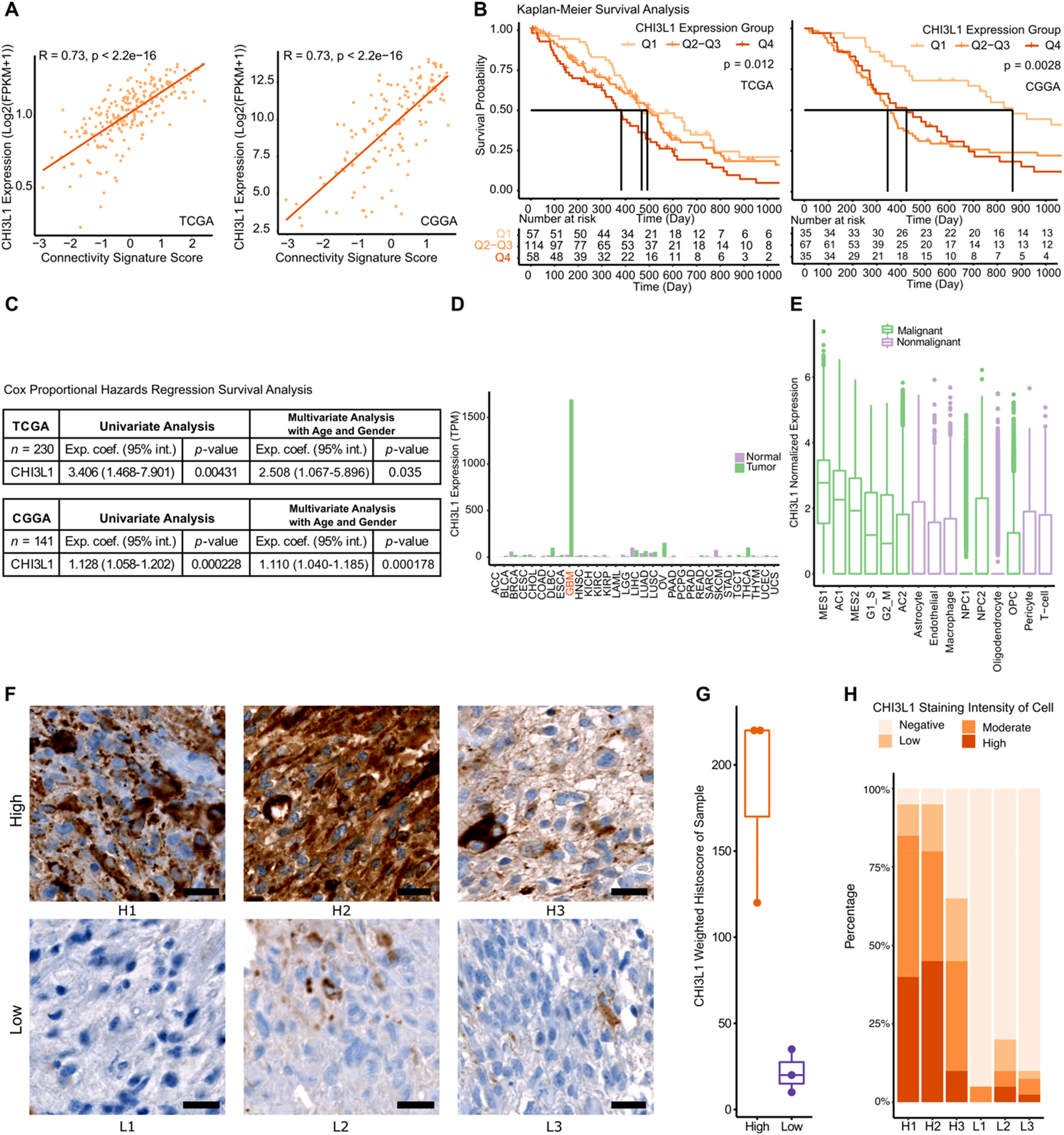
*CHI3L1* expression levels are correlated with connectivity. **A,** Scatter plot showing correlation between *CHI3L1* expression level (log2(FPKM+1)) and connectivity signature scores in 230 TCGA *IDH* wt GB RNA-Seq samples (left panel) and 141 CGGA *IDH* wt GB RNA-Seq samples (right panel). Connectivity signature scores were scaled and centered across samples, and winsorized to −3 and 3. Pearson correlation test was used to calculate correlation coefficients and p values. **B,** Kaplan-Meier survival analysis (Left, TCGA; Right, CGGA) according to *CHI3L1* expression groups (grouping by the first quartile [Q1], the two middle quartiles [Q2-Q3], and the last quartile [Q4] of *CHI3L1* expression levels). **C,** Cox proportional hazards regression survival analysis in cohorts (Top, TCGA; Bottom, CGGA). Univariate analysis with *CHI3L1* expression levels and multivariate analysis with *CHI3L1* expression levels (log2[FPKM+1]) adjusted for ages and genders. Exponents of the coefficients (Exp. coef.) with 95% confidence intervals (95% int.) indicated the hazard ratio of higher *CHI3L1* expression levels. **D,** Median *CHI3L1* expression levels (TPM) in 31 tumor types and related normal tissue retrieved from GEPIA. GB cohort is highlighted in red. **E,** Box plot of *CHI3L1* expression levels in malignant cell states and non-malignant cell types from snRNA-Seq dataset of 21 GB patient samples. **F-H,** Immunohistochemistry staining of CHI3L1 in three patients with high connectivity signature score (H1, H2 and H3), and three patients with low connectivity signature score (L1, L2 and L3). **F,** Representative images of CHI3L1 staining. **G,** Box plot of weighted histoscores of CHI3L1 staining per sample. **H,** Frequency of CHI3L1 staining intensity of cells per sample.

Together, this data suggests a functional role of CHI3L1 in tumor cell connectivity and CHI3L1 RNA and protein expression as an alternative way to determine overall tumor (cell) connectivity in GB if determination of the connectivity signature score by scRNA-Seq or RNA-Seq analysis is not possible.

### Higher connectivity is found in tumors of the mesenchymal expression subtype and *TP53* wt tumors

Next, the connectivity signature was applied to the 230 *IDH* wt GBs from the TCGA RNA-Seq dataset. The goal of this analysis was to identify the associations between the connectivity signature scores, expression subtypes (19), and gene mutations. The mesenchymal expression subtype was associated with the highest connectivity signature score, while the lowest score was observed in the proneural subtype **(Figure 6A)**. Mesenchymal subtype consisted mainly of MES1 and AC1 signatures, whereas classical tumors were found to be purely AC and proneural tumors had high frequency of OPC and NPC signatures **(Figure 6B)**. In the classical and proneural tumors, we found a higher fraction of the low connectivity AC2 signatures probably accounting for lower connectivity signature score **(Figures 6B-C)**. The associations of the connectivity signature score and expression subtypes were validated in the CGGA cohort (*n* = 141, **Supplementary Figure 8A-D**).

**Figure 6.**
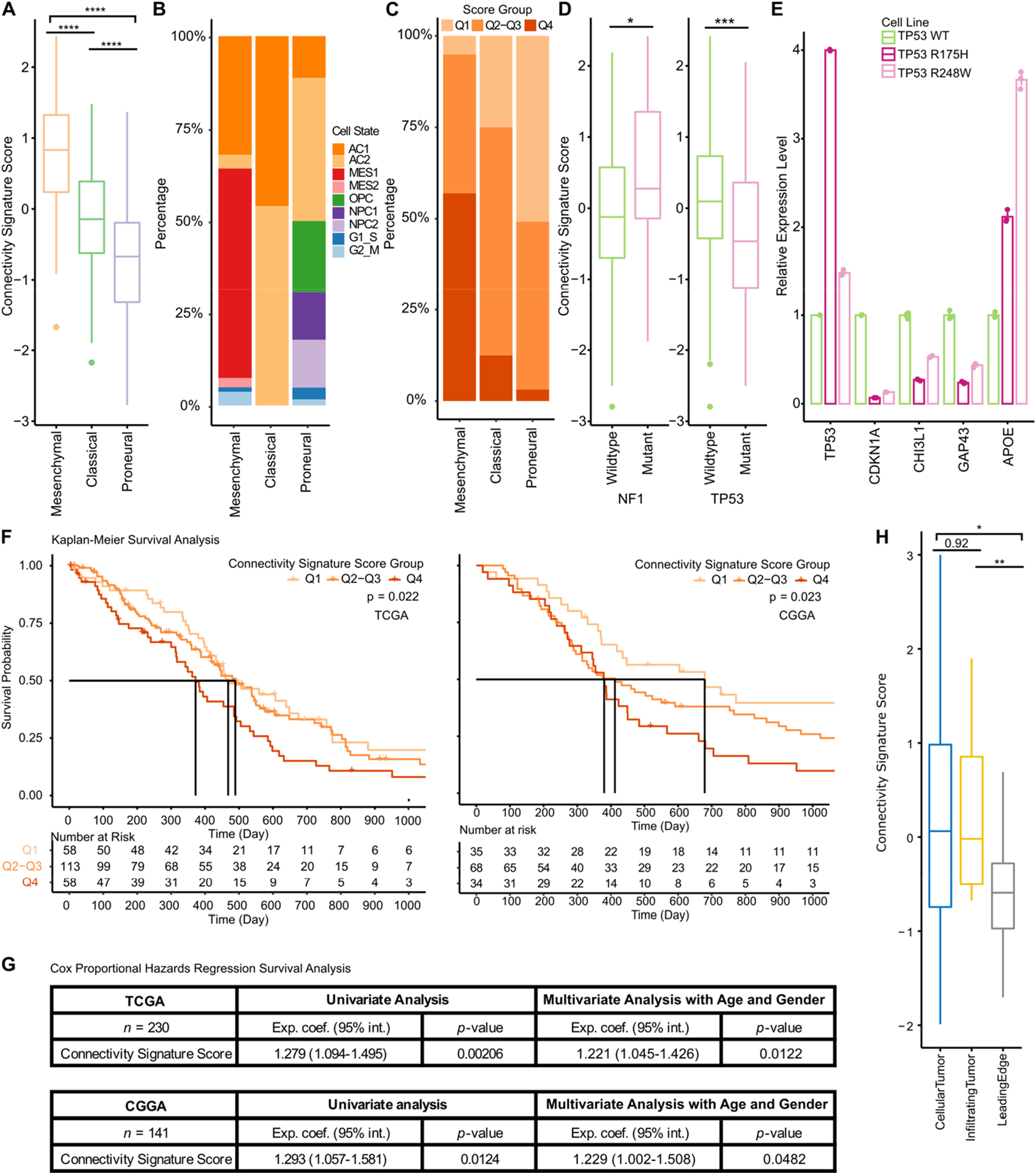
Expression subtype and patient prognosis are related to connectivity signature scores in validation cohorts. **A-D, F-G,** 230 TCGA GB samples. **F-G,** 141 CGGA GB samples. **H,** 73 IvyGAP GB samples. **A,** Box plot of connectivity signature scores in three GB expression subtypes (81 mesenchymal, 87 classical and 62 proneural). **B,** Frequency of dominant cell states in each expression subtype. **C,** Frequency of connectivity signature score groups in each expression subtype. Connectivity signature scores grouped by Q1, Q2-Q3 and Q4. **D,** Connectivity signature scores in samples grouped by mutation status (synonymous mutations were removed). Left, 195 *NF1* wt and 35 *NF1* mutated samples. Right, 173 *TP53* wt and 57 *TP53* mutated samples. P values were calculated by Mann-Whitney U test. *, p < 0.05; **, p < 0.01; ***, p < 0.001. **E,** Bar plot of relative gene expression by qPCR of *TP53, CDKN1A, CHI3L1, GAP43* and *APOE* in TP53-mutant overexpressing (*TP53 R175H* and *TP53 R248W*) against TP53 wt overexpressing (*TP53 WT*) GB cell lines (n = 3). **F,** Kaplan-Meier survival analysis in cohorts (Left, TCGA; Right, CGGA) stratified into groups using Q1, Q2-Q3 and Q4 of the connectivity signature score. **G,** Cox proportional hazards regression survival analysis in cohorts (Top, TCGA; Bottom, CGGA). Univariate analysis with connectivity signature scores and multivariate analysis with connectivity signature scores adjusted for ages and genders. **H,** Boxplot of connectivity signature scores in three structure groups (30 cellular tumor, 24 infiltrating tumor and 19 leading edge) from IvyGAP cohort. **A, D, H,** Connectivity signature scores were scaled and centered across samples per cohort, and winsorized to −3 and 3.

Among the recurrent mutated genes in at least 5% GB patients (27 genes), two mutated genes, neurofibromatosis type 1 (*NF1)* and tumor protein p53 (*TP53)*, were associated with connectivity signature scores (false discovery rate [FDR] < 0.25). *NF1* mutations were present in 35/230 (15%) of GB patients in the TCGA cohort, which are associated with the mesenchymal subtype (14), were correlated with higher connectivity signature scores. Even when comparing tumors only of the mesenchymal subtype, *NF1* mutations were still associated with a higher connectivity signature score (**Supplementary figure 8E**). *TP53* mutations were present in 57/230 (25%) of the GB patients in the TCGA cohort and were correlated with a lower connectivity signature score (**Figure 6D**), which might be in line with the *TP53* dependency of nanomembrane tube formation in astrocytes (20). Moreover, expression of *CHI3L1* was higher in *TP53* wt tumor tissue samples than in samples with *TP53* mutation (**Supplementary Figure 8F**). Despite the same trend, no significant difference was observed for *GAP43* and *APOE* (**Supplementary Figure 8F**). On a functional level overexpression of *TP53* wt in GB tumor cells had only a minor effect on *CHI3L1* expression (**Supplementary Figure 8G**). However, overexpression of the two different dominant mutant GB *TP53* hotspot mutations R175H and R248W inhibited *TP53* downstream activity as measured by with cyclin dependent kinase inhibitor 1A (*CDKN1A)* expression and most importantly reduced RNA expression of two main connectivity markers *CHI3L1* and *GAP43* (**Figure 6E**). This argues for a functional relation between *CHI3L1 and GAP43* expression and *TP53* mutations. Previous studies already suggested that functional TP53 is necessary for GAP43 expression and axon outgrowth (21), supporting the functional role of *TP53* mutations in tumor cell connectivity.

### Cell-to-cell connectivity is associated with worse patient survival

Importantly, the impact of tumor cell connectivity on patient survival remained unresolved so far. To clarify this point, multiple survival analyses of connectivity signature scores were performed in TCGA and CGGA *IDH* wt GB patient cohorts. The shortest survival was found for patients with the highest quartile of connectivity signature score (Kaplan-Meier survival analysis on three connectivity signature score groups, **Figure 6F**). A constant increase in the risk of death correlated with the increase of connectivity signature score (Cox proportional hazards regression survival analysis on continuous connectivity signature scores, **Figure 6G**). The association of the connectivity signature score with patient survival remained significant after adjusting for ages and genders in a multivariate analysis (**Figure 6G**).

Furthermore, we adjusted the survival analysis for ages, genders as well as expression subtypes in the TCGA cohort, and found that patients with higher connectivity signature scores had an increasing risk of death (p = 0.0128, **Supplementary Figure 9A**). To more specific, in proneural subtype patients, high connectivity signature score group had lower survival probability (p = 0.031, **Supplementary Figure 9B**). Mesenchymal subtype patients had a similar trend as proneural subtype patients, but not significant (p = 0.065, **Supplementary Figure 9B**). This trend was not found in classical subtype patients (**Supplementary Figure 9B**). As a comparison to connectivity signature score, we performed similar survival analysis for *CHI3L1* expression levels (**Supplementary Figure 9C-D**). *CHI3L1* expression did not show a significant association with patient survival after adjusting for expression subtypes (**Supplementary Figure 9C-D**). At this point of view, the 71-gene-synthesized connectivity signature score outperformed one-gene marker *CHI3L1*.

Even though the connectivity signature was not particularly developed for *IDH* mutant gliomas, the connectivity signature score proved to be higher in astrocytic (1p/19q intact) compared to oligodendroglial (1p/19q codeleted) *IDH* mutant gliomas, reflecting previous histology-based morphological data (6). In *IDH* mutant glioma, only a trend for worse survival (*p* = 0.097) was detectable for patients with high connectivity signature score (**Supplementary Figure 9E-G**).

Finally, as sampling of tumor tissue for sequencing is mainly performed in one spot per tumor in routine analysis, we estimated the impact of different locations in the tumor on the connectivity signature score by analyzing the Ivy Glioblastoma Atlas Project (IvyGAP) dataset. The connectivity signature scores were lower in the leading edge compared to cellular and infiltrating tumors (**Figure 6H**), in line with the known higher anatomical and functional tumor cell connectivity in more solid established glioma areas (6, 16).

## Discussion

While the discovery of communicating, self-repairing and resistant tumor cell networks has changed our understanding of incurable gliomas, with multiple clinical implications (22), the measurement of this crucial tumor cell connectivity in patient samples, and a deeper understanding of its molecular underpinnings remained elusive. In this study, a connectivity signature score was established that proved feasible and valid to rapidly assess the degree of TM connectivity in various gliomas, was associated with AC1 and MES1 cell states, the mesenchymal expression subtype, and with worse patient survival (**Figure 7**). Furthermore, a considerable number of known and unknown genes associated with TM connectivity in GB were identified. The unexpectedly high proportion of upregulated genes (34/40, 85%) in the scRNA-Seq-derived connectivity signature that have been previously associated with neurogenesis, neural tube development or glioma progression highlights its biological plausibility (6,22,23) and the utility of this new connectivity signature for further in-depth gene analysis and use in clinical studies.

**Figure 7.**
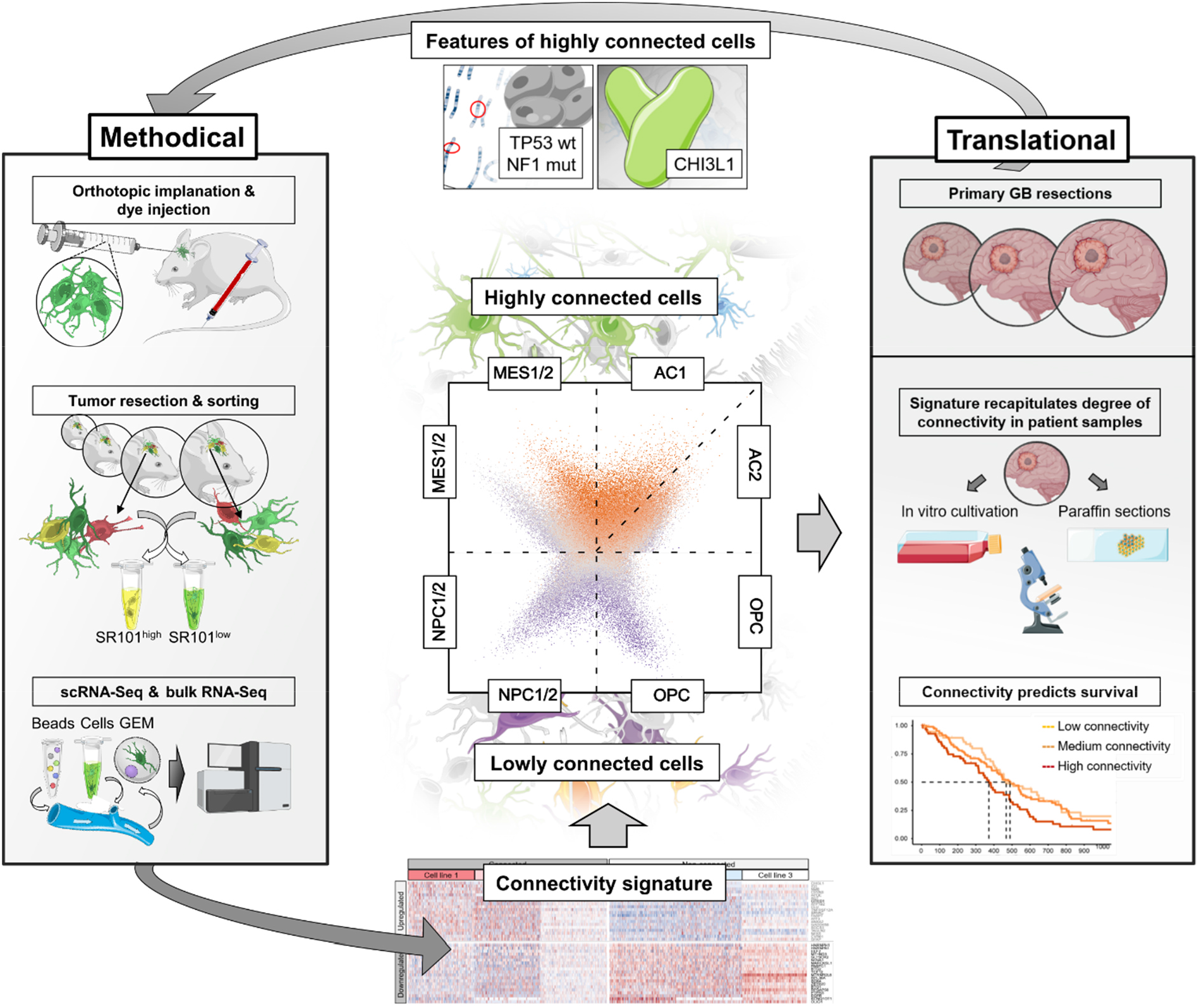
Summary and graphical abstract: TurboGFP-labeled PDGCLs (left box) were orthotopically implanted into mice. After tumor establishment, highly connected and lowly connected populations were sorted using a SR101 and sequenced by RNA-Seq and scRNA-Seq (left box). Differentially expressed genes revealed a potential molecular signature associated with connected and unconnected cells (bottom of the middle part). Applying this connectivity signature to previously reported GB cell states deciphered MES1 and AC as the ones with the highest connectivity signature score. Within the AC population, a subpopulation with an increased connectivity signature was identified (center). High-scoring cells were enriched for TP53 wt and NF1 mutation. CHI3L1 is a robust marker of highly connected cells (top box). *In vitro* PDGCL and patient paraffin sections with a high connectivity signature score showed a highly connected phenotype. Lastly, patients with a high connectivity signature score had a less favorable outcome than patients with a low score (right box).

For comparison to scRNA-Seq-derived connectivity signature, we also identified the RNA-Seq-derived connectivity signature. These two signatures have only a small proportion of overlapping genes (18% in scRNA-Seq-derived signature, 5% in RNA-Seq-derived signature), because of the differences between the sequencing technologies, especially the high drop-out rate in 10X scRNA-Seq technology. The correlation of fold changes in all 16,759 quantified genes between scRNA-Seq and RNA-Seq is low (R = 0.33, **Supplementary Figure 10A**). While focusing on the genes expressed in at least 10% cells of scRNA-Seq dataset, the correlation increased to medium (R = 0.56, **Supplementary Figure 10B**). While only focusing on the significant DEGs, the correlation further increased to R = 0.77 (**Supplementary Figure 10C**). The overlapping genes between two connectivity signatures have a high correlation (R = 0.89, **Figure 1H**). What’s more, the enriched GO biological processes, semantic similarity, and connectivity signature scores between two connectivity signatures have high concordances. These indicate a robustness of the development of connectivity signature, even with the two different experimental approaches.

It is challenging to measure tumor cell connectivity in patient samples robustly and reproducibly with standard histology (6). Therefore, a molecular gene expression connectivity signature that can be easily used to calculate a score has several advantages and can be applied on sequencing datasets from fresh tumor material, by RNA-Seq or – when available – scRNA-Seq.

Even simpler, *CHI3L1* expression correlates particularly well with the connectivity signature score, and the addition of recombinant CHI3L1 protein as well as antibody blocking modulates connectivity, strongly suggesting a functional role of CHI3L1 in tumor cell connectivity rather than a pure correlative marker. RNA expression or immunohistochemistry of CHI3L1 could therefore be a good estimator of tumor cell connectivity that could be easily used in a clinical setting from formalin-fixed paraffin embedded (FFPE) tissue when high-throughput scRNA-Seq or RNA-Seq methods are not feasible or available.

*TP53* mutations were identified to be associated with higher connectivity signature score. This is remarkable as besides *NF1* mutation as a mesenchymal marker no other small nuclear variant was associated with the connectivity signature score. *TP53* mutations may not be all alike and therefore more mutation type specific investigations are warranted in further studies.

Besides the increased TM-connectivity in AC tumor cells, which appears plausible given the principal connective nature of astrocytic cells, and AC tumor cells reflecting astrocytic programs, also a subset of MES tumor cells showed TM-connectivity. Accordingly, it has been observed long ago that “mesenchymal development” is characterized by single cells connected *via* long cellular processes to a functional syncytium” (24), not unlike the tumor cell networks in GB.

Finally, survival analysis showed a clear association of the tumors with the highest connectivity signature scores and worse patient outcome. This is in line with our previous preclinical findings that tumor cell connectivity by a cellular network of TM connections is a resistance factor to all standard glioma therapies (6,7,10,22). We have shown before that the prognostically favorable oligodendrogliomas (25, 26) do not form a relevant TM-connected tumor cell network and a low expression of key TM drivers. As a further validation, we confirmed a lower connectivity signature score in the oligodendroglial tumors (6, 16).

Besides O-6-methylguanine-DNA methyltransferase (*MGMT)* promoter methylation for alkylating chemotherapy, there is no clinically established marker for assessing the resistance to radiotherapy, chemotherapy, and surgery in glioma. The connectivity signature developed here can serve as a molecular fingerprint for a key mechanism of resistance, and thus as a prognostic and even predictive biomarker. This biomarker, however, needs further validation in prospective clinical trials before it can be used to assign patients to treatment groups in clinical trials. Finally, it can also serve as a biomarker for future disconnection strategies that are currently in preclinical development (27).

This proof-of-concept study to collectively investigate the genetic background of glioma cell-to-cell connectivity on single cell level has several limitations. The main limitation is based on the SR101 dye transfer model, which is a proven (11, 12) yet not dichotomic discriminator of existing or non-existing cellular connectivity in gliomas. Therefore, we implemented several confirmatory measures on functional experimental and anatomical levels to assure that changes in connectivity signature score are accompanied by true changes of TM network formation. Other limitations are related to the ability to transduce the connectivity signature from a xenografted patient derived model to patient samples. In particular, current practice of taking samples from one or at best a few parts of the tumor for RNA sequencing may not account for potential intratumoral heterogeneity of TM-connectivity which may relate to cell density, hypoxia, molecular heterogeneity and the microenvironment. Analyses of the IvyGAP shows lower connectivity results in the leading edge part of tumors, which at least argues for an important role of the side from where the sample is taken.

In conclusion, we developed a connectivity signature with a respective score calculation for GBs based on scRNA-Seq analysis and identified biologically plausible markers in highly connected tumor cells for further investigation and confirmation in preclinical and clinical trials. *CHI3L1* expression has emerged as the easiest to assess single marker gene of the signature that can even be determined in standard paraffin sections. This offers the opportunity to translate the recent fundamental insights into key elements of tumor biology in GB into clinical trials and ultimately into clinical practice.

## Acknowledgements

We thank Hrvoje Miletic for providing the P3XX and BG5 cell line. We thank the scOpen lab of the DKFZ, Heidelberg, Germany for providing support with the 10x Chromium Controller for scRNA-Seq. Sequencing capacity was kindly provided by the Genomics and Proteomics Core Facility of the DKFZ. We are grateful to the Omics Data Core Facility of the DKFZ for providing data storage capacity. We thank Katja Bauer and Birgit Kaiser, Laura Doerner, Lea Hofmann, Moritz Schalles and Hai-Yen Nguyen for expert technical support.

## Author contributions

Conceptualization: L.H., D.C.H., F.W., M.S., W.W., T.K.

Resources: R.W., M.L.S., C.H., F.S., F.W., M.S., W.W., T.K.

Data curation: L.H., D.C.H., T.K.

Software: L.H., M.L.S., M.S.,

Formal analysis: L.H., D.C.H, H.M., T.K.

Supervision: M.S., W.W., T.K.

Funding acquisition: F.W., M.S., W.W., T.K.

Validation: L.H., D.C.H, H.M., J.I., E.J., S.W., P.S., F.W., M.S., W.W., T.K.

Investigation: L.H., D.C.H, H.M., R.X., J.I., E.J., S.W., P.S., V.V., D.D.A., K.E., D.R., R.W., F.S., T.K.

Visualization: L.H., D.C.H.

Methodology: L.H., D.C.H., H.M., P.S., D.D.A., F.S., T.K.

Writing-original draft: L.H., D.C.H., T.K.

Project administration: W.W., T.K.

Writing-review and editing: all authors

## Methods

### Cell culture of PDGCL xenografted mouse models

PDGCLs S24 and T269 were established from freshly dissected GB tissue from adult patients after informed consent (28). PDGCLs P3XX and BG5 were kindly provided by Hrvoje Miletic, K. G. Jebsen Brain Tumour Research Centre, University of Bergen (29). All four tumors have been diagnosed as GB, *IDH* wt. Methylation profiling with the methylation EPIC array (#WG-317-1003, Illumina, San Diego, California, USA) was used to confirm GB origin. S24 is characterized by a GB receptor tyrosine kinase (RTK) I, whereas BG5, P3XX and T269 exhibit a GB RTK II methylation subtype (**Supplementary Table 1** and (30). PDGCLs were cultured as neurospheres under serum-free, non-adherent, “stem-like” conditions in PDGCL media, consisting of DMEM/F-12 (#11330-032, Life Technologies, part of ThermoFisherScientific, Waltham, Massachusetts, USA), B27 supplement (#17504044, Life Technologies, part of ThermoFisherScientific, Waltham, Massachusetts, USA), insulin (#I9278, Sigma, part of Merck, Darmstadt, Germany), heparin (#H4784, Sigma, part of Merck, Darmstadt, Germany) epidermal growth factor (EGF; #PHG0311, Life Technologies, part of ThermoFisherScientific, Waltham, Massachusetts, USA) fibroblast growth factor (FGF; #PHG0021, Life Technologies part from ThermoFisherScientific, Waltham, Massachusetts, USA).

In order to allow identification and re-isolation after tumor resection, PDGCLs were lentivirally transduced with the MISSION^®^ shRNA vector pLKO.1-puro-CMV-Turbo green fluorescent protein (TurboGFP)_shnon-target (#SHC016, Sigma, part of Merck, Darmstadt, Germany) for cytosolic TurboGFP expression. The production of lentiviral particles and cellular transductions was carried out as described previously (6). Successfully transduced tumor cells were selected using 1 µg/ml puromycin (#A2856.0100, Applichem, Darmstadt, Germany) and FACS sorting.

All four PDGCLs were regularly checked for authenticity and absence of infections, e.g. with mycoplasms and non-human cell contamination by multiplex cell contamination test (Multiplexion GmbH, Heidelberg, Germany).

### Correlation of SR101 staining with TMs in PDGCL xenografted mouse models

All *in vivo* experiments in this study were approved by the local authorities (Regierungspräsidium Karlsruhe, Germany) and compliant with the institutional laboratory animal research guidelines. All efforts were made to minimize animal suffering and to reduce the number of animals used according to the 3R’s principles. Experiments were carried out as already described (6). Striatal tumor bearing male nude mice (RRID:MGI:5653040, Charles River, Wilmington, Massachusetts, USA) were intravenously injected with SR101 (#S359, Invitrogen, part of ThermoFisherScientific, Waltham, Massachusetts, USA) dissolved in sterilized saline solution (#2350748, B. Braun Melsungen AG, Melsungen, Germany) using a dose of 0.12 mg per g body weight. Repetitive intravital 2-photon microscopy was performed after SR101 injection using a Zeiss 7MP microscope (Zeiss, Oberkochen, Germany) equipped with a Coherent Chameleon UltraII laser (Coherent, Santa Clara, California, USA) and a band-pass 500–550 and 575–610 nm filter. SR101 was excited at 900 nm and TurboGFP at 950 nm 38-150 cells were analyzed in n=5 regions in 3 animals on D64+/-9 days. SR101 intensities of highly connected and lowly connected GB cells were measured in the cell bodies. The signal intensity was normalized by the mean value of the highest 10% of intensities in the respective region.

### Separation of highly and lowly connected cells in PDGCL xenografted mouse models

PDGCL spheroids were dissociated into a single cell suspension using Stem-Pro Accutase™ (#1110501, ThermoFisherScientific, Waltham, Massachusetts, USA). 5×10^4^ viable cells were slowly injected into the right hemisphere of 8-10 week old male nude mice (RRID:MGI:5653040; Charles River, Wilmington, Massachusetts, USA) using a 10 µl micro-syringe (#80308, Hamilton, Reno, Nevada, USA) driven by a stereotactic device (Stoelting, Wood Dale, Illinois, USA). The exact coordinates were 2 mm right lateral of the bregma and 1 mm anterior to the coronal suture with an injection depth of 2-3 mm below the dural surface. Tumors were grown until the mice showed first symptoms or ≥20% weight loss were met. Mice were intraperitoneally injected with SR101 (#S359, Invitrogen, S359, Invitrogen, part of ThermoFisherScientific, Waltham, Massachusetts, USA) dissolved in sterilized saline solution (#2350748, B. Braun Melsungen AG, Melsungen, Germany) using a dose of 0.12 mg per g body weight. After an incubation period to ensure maximum SR101 uptake from PDGCL cells, mice were deeply anesthetized with ketamine/Ketaset^®^ (#794-523, Zoetis, Berlin, Germany) and xylazine/Rompun^®^ (#770-081, Bayer, Leverkusen, Germany) and transcardially perfused with sterilized phosphate buffer saline (PBS, #D8537, Sigma, part of Merck, Darmstadt, Germany). The whole brain samples were removed and prepared into cell suspension using brain tumor dissociation kit (#130-095-942, Miltenyi Biotec, Bergisch Gladbach, Germany) and gentleMACSTM Dissociator (#130-093-235, Miltenyi Biotec, Bergisch Gladbach, Germany). The cell pellet was resuspended in FACS buffer, consisting of 1% fetal calf serum (FCS; #S0615, Sigma, part of Merck, Darmstadt, Germany) in PBS, and proceeded with FACS sorting.

#### FACS

The single cell suspension freshly prepared from xenografted brains was incubated with eBioscience™ Calcein Violet 450 AM (#65-0854-39, Invitrogen, part of ThermoFisherScientific, Waltham, Massachusetts, USA) and TO-PRO™-3 Iodide (#T3605, Invitrogen, part of ThermoFisherScientific, Waltham, Massachusetts, USA) for 10 min on ice prior to sorting. Standard gating techniques were used to discriminate doublets and dead cells. The viable fraction was defined by TO-PRO™-3 Iodide negativity and Calcein Violet 450 AM positivity. To further allow discrimination of the non-malignant cells, the TurboGFP population was selected for separation of highly connected tumor cells (SR101^high^) and lowly connected tumor cells (SR101^low^) using the FACSAria™ cell sorter (BD Biosystems, Franklin Lakes, New Jersey, USA). The following filters were used: V450/50 (Calcein Violet), B530/30(TurboGFP), YG586/15 (SR101) and R650/17 (TO-PRO™-3).

### RNA-Seq data generation and preprocessing from PDGCL xenografted mouse models

Sorted tumor cells were resuspended in lysis buffer included as a part of the RNeasy^®^ Micro Kit (#74004, Qiagen, Hilden, Germany). mRNA was then isolated and purified in accordance with the manufacturer’s instructions. The conversion of RNA to DNA was done with the SMARTer^®^ Ultra^®^ Low Input RNA for Illumina Sequencing (#634940, TakaraBio, Kusatsu, Japan). The libraries were then prepared using NEBNext^®^ ChIP-Seq Library Prep Master Mix Set for Illumina (#E6240, New England Biolabs, Ipswich, Massachusetts, USA) and sequenced on an Illumina HiSeq 2000 sequencer (RRID:SCR_020132, v.4, Illumina, San Diego, California, USA) in 50 bp single-end mode by Genomics and Proteomics Core facility, DKFZ. The bioinformatics tools for gene expression quantification from RNA-Seq were used with default parameters: The quality of bases was evaluated and controlled using FASTX-Toolkit (RRID:SCR_005534). HOMER (RRID:SCR_010881, v.4.7) was applied for PolyA-tail trimming; reads with a length of < 17 bp were removed. The filtered reads were mapped with STAR (RRID:SCR_004463, v.2.3) against the human reference genome (GRCh38) and Picard (RRID:SCR_006525, v.1.78) with CollectRNASeqMetrics were used for quality checking. Count data were generated by htseq-count (RRID:SCR_011867, v.0.9.1) using the GENCODE (RRID:SCR_014966, v26) for annotation. Genes with a total count of less than 10 were discarded.

### scRNA-Seq data generation from PDGCL xenografted mouse models

A total of 5×10^4^ highly and lowly connected cells from at least 3 mice/replicates per PDGCL suspension were FACS-sorted and subjected to a 10x Chromium Controller (10x Genomics, Pleasanton, California, USA) and further processed according to the manufacturer’s instructions. The technology samples a pool of around 750,000 barcodes to separately index each cells transcriptome. In brief, 10x barcoded gel beads are mixed with cells, enzyme and partitioning oil to create single cell gel beads in emulsion. Barcoded cDNA is generated by reverse transcription so that cDNA from individual cells share a common barcode. Afterwards, sequencing was carried out on a HiSeq 4000 sequencer (SY-401-4001, Illumina, San Diego, California, USA) or on a NovaSeq 6000 sequencer (20012850, Illumina, San Diego, California, USA) to obtain approximately 2 x 350 million reads per sample.

### Single nuclei (sn)RNA-Seq data generation from patient samples

#### Case selection and ethics approval

All 21 patients included have been treated at the Heidelberg University Hospital. All patients gave informed consent either prior to inclusion to the NCT Neuro Master Match (N²M²) pilot study (31) or to exploratory molecular analyses. The research is conducted in concordance with the declaration of Helsinki and was approved by the Ethics Committee at the University of Heidelberg, Germany (applications 206/2005 and AFmu-207/2017). The N²M² pilot study included patients with *MGMT* promoter unmethylated tumors, leading to an enrichment of *MGMT* promoter unmethylated samples in our analysis (18/21, 86%).

Frozen resected tumor material was retrieved from the Department of Neuropathology in Heidelberg and reviewed by a board-certified neuropathologist. Diagnoses were molecularly confirmed according to the recent WHO classification and methylation profiles were confirmed with methylation EPIC array (#WG-317-1003, Illumina, San Diego, California, USA).

For single nuclei isolation, resected tumor material underwent the following quality control. Exclusively material with a tumor content ≥ 70% and a low percentage of necrosis, as determined on hematoxylin and eosin-stained sections by a board-certified neuropathologist (Department of Neuropathology, University Hospital Heidelberg, Germany) was considered for further processing. Clinical and pathological characterization of patients are summarized in Table 1. Human patient samples were anonymized manually.

#### Single nuclei preparation

Tumor sections were roughly chopped on ice and resuspended in lysis buffer consisting of 320 mM sucrose (#84097, Sigma, part of Merck, Darmstadt, Germany), 5 mM CaCl2 (#21115, Sigma, part of Merck, Darmstadt, Germany), 3 mM Mg acetate (#63052, Sigma, part of Merck, Darmstadt, Germany), 2 mM EDTA (#AM9260G, Invitrogen, part of ThermoFisherScientific, Waltham, Massachusetts, USA), 0.5 mM ethylene glycol tetraacetic acid (EGTA, #J61721, Alfa Aesar, part of ThermoFisherScientific, Waltham, Massachusetts, USA), 1 mM dithiothreitol (DTT; #43816, Sigma, part of Merck, Darmstadt, Germany), 0.1% Triton X-100 (#A4975, AppliChem, Darmstadt, Germany) and 10 mM Tris(hydroxymethyl)aminomethan (Tris) pH 8.0 (#15568025, Life Technologies, part of Thermo Fisher Scientific, Waltham, Massachusetts, USA). The suspension was transferred to a dounce homogenizer (#9651617, Th. Geyer, Renningen, Germany) for nuclei isolation. Large debris was removed by 100 µm (#542000, Greiner Bio-one, Kremsmünster, Austria) and 70 µm (#542070, Greiner Bio-one, Kremsmünster, Austria) strainer meshes and the suspension collected in separate 50 ml tubes (#227261, Greiner Bio-one, Kremsm ü nster, Austria). Next, nuclei were subjected to three repeated wash cycles consisting of centrifugation (550 g, 5 min, 4°C), supernatant removal and resuspension in 1.5 ml washing buffer. Adaptions for the last cycle included addition of 500 µl homogenization buffer (320 mM Sucrose, 30 mM CaCl2, 18 mM Mg(Ac)2, 0.1 mM EDTA, 0.1% Nonidet P40 [#APA1694.0250, Applichem, Darmstadt, Germany], 0.1 mM phenylmethylsulfonyl fluoride [PMSF, #6367.2, Roth, Karlsruhe, Germany], 1 mM beta-Mercaptoethanol [#M7522, Sigma, part of Merck, Darmstadt, Germany], 60 mM Tris pH 8.0) to the nuclei pellet and a resting time of 5 min before resuspension in another 1 ml homogenization buffer. Further purification was done using a iodixanol (#07820, Stem Cell Technologies, Vancouver, Canada) gradient. Briefly, the pellet was resuspended in 200 µl gradient buffer consisting of 30 mM CaCl2, 18 mM Mg(Ac)2, 0.1 mM PMSF, 1 mM beta-Mercaptoethanol and 60 mM Tris pH 8.0. After transfer to a new microcentrifuge tube, 200 µl of 50% iodixanol in gradient buffer was used to generate a final concentration of 25% iodixanol. The nuclei suspension was carefully layered onto a gradient consisting of equivoluminous 300 µl layers of 29% and 35% iodixanol in gradient buffer supplemented with 160 mM sucrose. Separation was performed at 4°C for 20 min with 3000 g. 200 µl of the nuclei-containing interphase was collected and passed through a 20 µm filter (#130-101-812, Miltenyi Biotec, Bergisch Gladbach, Germany). Partially, trituration using wide-bore tips (#10089010, Thermo Fisher Scientific, Waltham, Massachusetts, USA) was necessary to facilitate disaggregation of the nuclei.

All aforementioned steps were performed on ice and all plastic consumables having contact with nuclei were pre-coated with 0.1% Triton X-100 prior to use to prevent sample loss.

Finally, integrity and purity of the nuclei was confirmed using Trypan Blue (#15250-061, LifeTechnologies, part of Thermo Fisher Scientific, Waltham, Massachusetts, USA) staining and the nuclei sequenced according to the 10x protocol (see section “scRNA-Seq data generation from PDGCL xenografted model”).

### Single cell data preprocessing

The gene expression count matrices of PDGCL xenografted mouse models scRNA-Seq were generated using Cell Ranger (RRID:SCR_017344, v.2.1.1, 10X Genomics) with default parameters, against the pre-built hg19 human reference genome (Cell Ranger reference, v.1.2.0). The count matrices of patient samples snRNA-Seq were generated using Cell Ranger (RRID:SCR_017344, v.3.0.1, 10X Genomics) with standard parameters, against a custom pre-mRNA hg19 human reference genome generated by mkref function following the official guideline (https://support.10xgenomics.com/single-cell-gene-expression/software/pipelines/3.1/advanced/references). We discarded cells by uniform exclusion criteria: (1) discarding cells which had fewer than 200 or more than 8,000 genes detected. (2) discarding cells which had fewer than 500 or more than 80,000 counts detected. (3) discarding cells which had a percentage of counts that came from mitochondrial genes of more than 10%.

After the uniform exclusion, sample-wise outlier cells were detected and removed if the number of genes or counts are more than three median absolute deviations (MADs) above the median using isOutliers function in the scater (RRID:SCR_015954, v.1.10.1). In each sample, per-cell doublet scores and per-sample doublet score thresholds were estimated by Scrublet (RRID:SCR_018098, v.0.2.1) with default parameters. If one doublet score threshold was located between two peaks of a doublet score histogram, this threshold was accepted and the cells with a doublet score higher than this threshold were removed. Unsupervised clusters were visualized in uniform manifold approximation and projection (UMAP) to further detect low quality clusters using Seurat (RRID:SCR_007322, v.3.1.5). In PDGCL xenografted mouse model dataset, outlier clusters were removed according to the MADs of the median number of genes or embeddings of UMAP in clusters. In patient samples dataset, one cluster that expressed markers of two different cell types was removed.

In the end, we obtained 35,822 cells from six samples of three PDGCL xenografted mouse models and 213,444 cells from 21 patient samples.

### Single cell data processing and integration

#### Data processing

After data preprocessing and quality control, scRNA-Seq data of PDGCL xenografted mouse models and patient samples were further processed using Seurat (RRID:SCR_007322, v.3.1.5) with default parameters: The gene expression counts were normalized using the NormalizeData function. Then 2000 highly variable genes were identified using the FindVariableFeatures function. The variation of number of counts among cells was regressed out, and the resulting residuals were scaled and centered by the ScaleData function. Next, we reduced dimensionality of the data by principal component analysis using the RunPCA function. The number of principal components (PCs) used for further analyses was determined using the ElbowPlot function (PDGCL dataset: 11 PCs; patient dataset: 24 PCs). The data was visualized in UMAP using RunUMAP function with determined PCs.

#### Data integration

To remove the differences of individuals and perform batch correction, an integration method based on identification of shared ‘anchors’ between pairs of samples was applied using the Seurat (RRID:SCR_007322, v.3.1.5) with default parameters: The gene expression count of each PDGCL or patient sample was normalized and selected highly variable genes using the NormalizeData and FindVariableFeatures functions. Then the normalized data (three PDGCLs or 21 patient samples) were integrated with the FindIntegrationAnchors function (dims = 1:30) and the IntegrateData function (dims = 1:30). The integrated data was used the ScaleData, RunPCA, ElbowPlot, RunUMAP functions as section “*Single cell data processing and visualization*” (PDGCL integrated dataset: 24 PCs; patient integrated dataset: 22 PCs).

### Identification of malignant and non-malignant cell types in snRNA-Seq of patient samples

#### Cell type signature scores

In patient integrated snRNA-Seq dataset, cell type signature scores (i.e., malignant signature score, macrophage signature score, T-cell signature score, oligodendrocyte signature score, endothelial signature score, pericyte signature score, and astrocyte signature score) based on cell type markers (see the next paragraph) were calculated in each cell using the AddModuleScore function in Seurat (RRID:SCR_007322, v.3.1.5).

#### Cell type marker collections

The top 100 upregulated markers per cell types (i.e., malignant cells, macrophages, T-cells and oligodendrocytes) were identified from a GB scRNA-Seq dataset (5) using the FindAllMarkers function with default parameters in Seurat (RRID:SCR_007322, v.3.1.5). The top 100 upregulated markers of endothelial cells were obtained from a healthy brain RNA-Seq dataset (17).The top 100 enriched markers in pericytes were obtained from brain mural cells RNA-Seq dataset (18). The upregulated markers of healthy astrocytes compared to malignant astrocytes were obtained from a human brain RNA-Seq dataset (17).

#### Cell type assignment

The patient integrated dataset was performed unsupervised clustering using the FindNeighbors function with 22 PCs and the FindClusters function (resolution = 0.7), 24 clusters were obtained. In each cluster, the medians of each cell type signature score were calculated and represented as *S_ij_*, with *i* being one cell type and *j* being one cluster. Then the non-malignant scores *NMS_ij_* were defined as *S_ij_* minus malignant signature score *S_mj_* (m indicates malignant cells): *NMS_ij_ = S_ij_ − S_mj_*. The clusters were assigned to non-malignant cell types if *NMS_ij_* more than MAD above the median of all *NMS_ij_* : cluster 8, 9, and 23 as macrophages, cluster 5 as oligodendrocytes, cluster 19 as T-cells, cluster 22 as pericytes and cluster 17 as endothelial cells. The remaining clusters were assigned as malignant clusters and were validated based on CNV estimation using the infercnv (RRID:SCR_021140, v.1.2.1) with recommended parameters for 10x Genomics data (cutoff = 0.1, cluster_by_groups = TRUE, denoise = TRUE, HMM = TRUE). The assigned macrophages, oligodendrocytes, T-cells, pericytes and endothelial cells were used as reference non-malignant cells. Each non-malignant cell type and malignant clusters were downsampled to 500 cells. We found that the malignant clusters contained large-scale CNVs except cluster 21. The cluster 21 showed the highest astrocyte signature score and, accordingly, cluster 21 was reassigned as astrocyte cluster.

### Development of the connectivity signatures

In scRNA-Seq data of the PDGCL xenografted models, DEGs between highly and lowly connected groups were identified in each PDGCL xenografted model using the FindMarkers function with default parameters in Seurat (RRID:SCR_007322, v.3.1.5). We then aggregated the significant DEGs (adjusted p value < 0.05) from all three PDGCLs. Among the aggregated DEGs, we examined the direction of regulation of the DEGs, only the DEGs which were significantly differentially expressed with the same direction of regulation in at least two PDGCLs were kept. The remaining DEGs were further refined to obtain strongly regulated genes with an absolute log fold-change greater than 0.4. 50 DEGs were obtained. Additionally, the FindConservedMarkers function with default parameters was used to identify conserved DEGs between highly and lowly connected groups irrespective to PDGCLs. Among the conserved DEGs, the DEGs regulated in the same direction across all three PDGCL xenografted models were kept. 21 additional DEGs were obtained. In total, 71 DEGs were derived from scRNA-Seq dataset and served as a connectivity signature.

In RNA-Seq of PDGCL xenografted models, DEGs between highly and lowly connected groups were identified using DESeq2 (RRID:SCR_015687, v.1.22.2): The PDGCL xenografted models information was included in the design formula of the DESeqDataSet function to obtain conserved DEG of highly and lowly connected groups across PDGCL xenografted models. Differential expression analysis was performed using the DESeq function. Then the results were shrinked with apeglm method in the lfcShrink function. Other parameters are by default. The significant DEGs (adjusted p value < 0.05) with an absolute log2 fold-change greater than 1 were kept. Finally, 245 DEGs were derived from the RNA-Seq dataset.

### Heatmap visualization of the connectivity signatures

For each connectivity gene derived from scRNA-Seq, the gene expression level of the gene in cells of each sample were averaged using the AverageExpression function in Seurat (RRID:SCR_007322, v.3.1.5). The average expression levels were scaled, centered, winsorized at −3 and 3, and then visualized as heatmap using ComplexHeatmap (RRID:SCR_017270, v.2.5.4).

The bulk count matrix was transformed with variance stabilizing transformation using the vst function in DESeq2 (RRID:SCR_015687, v.1.22.2), and the batch effects between the PDGCL xenografted models were corrected with the removeBatchEffect function of the LIMMA package (RRID:SCR_010943, v.3.36.5). Finally, the expression levels of connectivity genes derived from RNA-Seq were scaled, centered, winsorized at −3 and 3, and then visualized as heatmap using ComplexHeatmap (RRID:SCR_017270, v.2.5.4).

### GO enrichment analysis

GO enrichment analysis of connectivity signature derived from scRNA-Seq (71 genes) or connectivity signature derived from RNA-Seq (245 genes) was performed by the compareCluster function using clusterProfiler (RRID:SCR_016884, v.3.18.1) against “GO Biological Process” with setting *fun = enrichGO* and *ont = “BP”*. The most enriched GOs were visualized with the emapplot function using enrichplot (v.1.10.2). The semantic similarity of both connectivity signatures against GO biological process were performed by mclusterSim function using GOSemSim (v.2.16.1).

There are 16,759 genes commonly expressed in both scRNA-seq and RNA-Seq datasets of PDGCL xenografted models. Gene set enrichment analysis of these genes preranked by the fold change between highly and lowly connected groups in the scRNA-Seq dataset or the fold change between highly and lowly connected groups in the RNA-Seq dataset was calculated by Gene Set Enrichment Analysis (RRID:SCR_003199, v.4.1.0) against “neurogenesis” gene set.

GO enrichment analysis of 100 DEGs between the two AC subgroups was performed using clusterProfiler (RRID:SCR_016884, v.3.18.1) against GO biological process terms (Molecular Signatures Database, RRID:SCR_016863, v.7.1).

### Connectivity signature score

The connectivity signature derived from scRNA-Seq data contains 71 genes, among which, 40 genes are upregulated in highly connected cells and 31 genes are downregulated. The 40 upregulated genes were used as a gene set to calculate a score (connectivity-upregulated signature score) in each cell using the AddModuleScore function in Seurat (RRID:SCR_007322, v.3.1.5). The score represents the relative expression of a gene set. Similarly, a second score (connectivity-downregulated signature score) based on the 31 downregulated genes was calculated. Finally, the connectivity signature score was defined as the connectivity-upregulated signature score minus the connectivity-downregulated signature score. Another connectivity signature score based on 245 genes (57 upregulated genes and 188 downregulated genes) derived from the RNA-Seq data were generated accordingly.

### The performance of the connectivity signatures for prediction of SR101-sorted labels

In each cell of scRNA-Seq data from PDGCL xenografted models, connectivity-upregulated signature score based on 40 scRNA-Seq-derived upregulated connectivity genes and connectivity-downregulated signature score based on 31 scRNA-Seq-derived downregulated connectivity genes were calculated. If the connectivity-upregulated signature score was higher than the connectivity-downregulated signature score, the cell was predicted as “Highly connected” cell, otherwise, the cell was predicted as “Lowly connected” cell. Confusion matrix and prediction metrics (i.e accuracy, sensitivity, specificity, positive predictive value and negative predictive value) were obtained between the number of cells predicted as “Highly connected” or “Lowly connected” base on calculated scores and the number of cells labelled as “Highly connected” or “Lowly connected” after SR101-based cell sorting, using R package caret (RRID:SCR_021138, v.6.0-80). Another prediction based on 57 RNA-Seq-derived upregulated connectivity genes and 188 RNA-Seq-derived downregulated connectivity genes were calculated in the same way.

#### Negative control

100 random gene sets, each gene set including 71 randomly selected genes (40 gene as an upregulated gene set and 31 as a downregulated gene set, the same as scRNA-Seq-derived connectivity signature), were utilized to calculate scores and obtained the average prediction metrics. Another 100 random gene sets, each gene set including 245 randomly selected genes (57 gene as an upregulated gene set and 188 as a downregulated gene set, the same as RNA-Seq-derived connectivity signature), were utilized to calculate scores and obtained the average prediction metrics.

### Malignant cell state assignment

Cell state markers from a GB scRNA-Seq study (5) were utilized to calculate cell state signature scores in each malignant cell in our patient sample snRNA-Seq dataset using the AddModuleScore function in Seurat (RRID:SCR_007322, v.3.1.5). Malignant cells were assigned to this cell state that gained the highest signature score among all cell state signature scores.

### Two-dimensional projection of patient malignant cells by cell state

Similar to (5), we obtained signature scores for each cell state in single cells and projected the cells according to the cell state signature scores. Y axis values represent the maximum score from the AC/MES1/MES2 states from which the maximum score from the OPC/NPC1/NPC2 states have been subtracted. If Y *>* 0, the X axis values represent AC minus the maximum of MES1 and MES2. If Y ≤ 0, the X axis values represent OPC minus the maximum of NPC1 and NPC2. Cells were colored by connectivity scores and plotted by ggplot2 (RRID:SCR_014601, v.3.3.2).

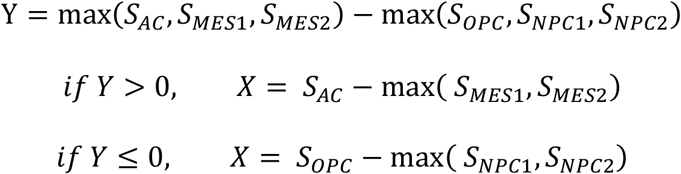

### Identification of subpopulations of astrocyte-like cells

In the two-dimensional projection of astrocyte-like cells only, we separated cells into two groups by a line with slope 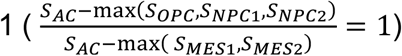. We obtained one group with higher connectivity scores and the other with lower connectivity scores. DEGs between these two groups were identified using the FindMarkers function in Seurat (RRID:SCR_007322, v.3.1.5).

### PDGCL in vitro models of connectivity

#### Quantification of TMs

PDGCLs S24, BG5, P3XX and T269 stably transduced with cytosolic TurboGFP, as previously described (6, 16), were used to allow visualization of TMs during confocal microscopy. All PDGCLs were cultured under two different culture conditions. For TM-, non-adherent conditions cells were cultured in PDGCL media as described before. In order to induce the formation of TMs, cells were kept in adherent culture conditions using DMEM (#11965-118, Life Technologies, part of ThermoFisherScientific, Waltham, Massachusetts, USA) supplemented with 10% FCS. Briefly, ethidium-homodimer 2 (EthD2, #E3599, Invitrogen, part of ThermoFisherScientific, Waltham, Massachusetts, USA) and Hoechst33342 (#H3570, Invitrogen, part of ThermoFisherScientific, Waltham, Massachusetts, USA) were added. Images were acquired on a LSM710 confocal microscope (Zeiss, Oberkochen, Germany) and an EC plan Neofluar^®^ 10×0.3 M27 objective (Zeiss, Oberkochen, Germany). The following excitations and detection wavelengths were used: 405/410-587 (Hoechst33342), 488/493-598 (TurboGFP) and 561/597-685 (EthD2). Laser power and maximum imaging time were tuned as low as possible to avoid phototoxicity. Images with a pixel size of 0.89 µm and an imaging frequency of 0.3 Hz were used for quantification. Mann-Whitney U test was used to compare connectivity in TM+ and TM-conditions.

#### Image processing and quantification

Multi photon laser scanning microscope (MPLSM) data were acquired by Zeiss ZEN^®^ Black Software (RRID:SCR_018163, Zeiss, Jena, Germany), which was also used for primary image calculation. Images were then transferred to Fiji 2.0.0 (RRID:SCR_002285) for analysis and processing. In the figures, maximum intensity projections are shown. Analyses were performed semi-automatically using the pixel classification and object quantification workflows in Ilastik software (RRID:SCR_015246, DOI:10.1109/ISBI.2011.5872394) after appropriate training.

### scRNA-Seq data generation from PDGCL in vitro models of connectivity

#### Cell lines and cell culture

PDGCLs S24, BG5, P3XX, T269 were cultured under TM- and TM+ conditions as described before.

#### FACS

After cultivation under both culture conditions, cells were blocked with 1% BSA in PBS. Cells were washed with PBS and subsequently resuspended in 1.5 ml of PBS/1%BSA containing 1 µM calcein AM (#C1430, Life Technologies, part of ThermoFisherScientific, Waltham, Massachusetts, USA) and 0.33 µM TO-PRO™-3 (#T3605, Invitrogen, part of ThermoFisherScientific, Waltham, Massachusetts, USA) to co-stain before sorting. Sorting was performed with FACSAria™ Fusion Special Order System (BD Biosystems, Franklin Lakes, New Jersey, USA) using 488nm (Calcein AM, 530/30 filter) and 640nm (TO-PRO-3™, 670/14 filter) lasers. An unstained control was included with every sample. Standard, strict forward scatter height versus area criteria were applied to discriminate doublets and gate only for single cells. Viable cells were detected as staining positive for calcein AM and negative for TO-PRO™-3.

#### scRNA-Seq

Cells were sorted into 96 well plates (#0030128.648, Eppendorf, Hamburg, Germany) containing cold TCL Buffer (#1070498, Qiagen, part of ThermoFisherScientific, Waltham, Massachusetts, USA) including 1% beta-mercaptoethanol (#M7522, Sigma, part of Merck, Darmstadt, Germany), snap frozen on dry ice and stored at −80°C. Whole transcriptome amplification, library preparation and sequencing were performed according to the SmartSeq2 protocol (32) with the following modifications as previously published (5): RNA purification from single cells was performed with Agencourt RNAClean XP beads (#A63987, Beckmann Coulter, Brea, California, USA) prior to olio-dT primed reverse transcription with Maxima reverse transcriptase (#EP0753, Life Technologies part of Thermo Fisher Scientific, Waltham, Massachusetts, USA) and locked template switch oligonucleotide (#339413, Qiagen, part of Thermo Fisher Scientific, Waltham, Massachusetts, USA). This was followed by 20 cycles of polymerase chain reaction (PCR) amplification using KAPA HiFi HotStart ReadyMix (#KK2602, Roche, Basel, Switzerland) and subsequent purification with Agencourt AMPure XP beads as described. Library construction was performed using the Nextera XT Library Prep kit (#FC-131-1024, Illumina, San Diego, California, USA) and custom barcode adapters (sequences available upon request). Libraries from 864 cells with unique barcodes were combined and sequenced with a NextSeq 500 sequencer (#SY-415-1001, Illumina, San Diego, California, USA).

### scRNA-Seq data processing of PDGCL in vitro models of connectivity

Sequencing reads were aligned using STAR (RRID:SCR_004463, v.2.5.3a) against the human reference genome hg19, and gene counts were generated and annotated using GENCODE (RRID:SCR_014966, v19) by featureCounts function of Subread package (RRID:SCR_009803, v.1.5.3). Gene counts were normalized to fragments per kilobase million (FPKM) values and log2 transformed. We identified low quality cells by the number of expressed genes lower than 2000 or higher than 8000. We obtained 735 cells from four PDCGLs. Then the data were integrated by ‘anchors’ and visualized in UMAPs with Seurat (RRID:SCR_007322, v.3.1.5).

### Quantitative real-time polymerase chain reaction (qPCR)

#### RNA extraction and cDNA synthesis

Harvested cells were washed with ice-cold PBS (#D8537-500ML, Sigma, part of Merck, Darmstadt, Germany). Afterwards, cells were resuspended in 1% beta-Mercaptoethanol (#M3148-100ml, Sigma, part of Merck, Darmstadt, Germany)-supplemented RLT lysis buffer, which is part of the QIAGEN RNeasy MicroKit (#79216, Qiagen, Hilden, Germany) or QIAGEN RNeasy Mini Kit (#74004, Qiagen, Hilden, Germany).

The kit type for subsequent RNA extraction was tailored to the absolute cell numbers. Lysates containing up to 500,000 cells were processed with the QIAGEN RNeasy^®^ Micro Kit whereas samples with 500,000 to one million cells were processed with the QIAGEN RNeasy^®^ Mini Kit. All steps were carried out according to the manual. On column DNAse digestion was performed with the RNAse free DNAse set (#79254, Qiagen, Hilden, Germany). RNA was eluted into RNAse-free water (#4387936, ThermoFisherScientific, Waltham, Massachusetts, USA). Reverse transcription was performed according to the manufactureŕs recommendations using the High-Capacity cDNA Reverse Transcription Kit with RNAse Inhibitor (#4374967, Applied Biosciences Applied Biosciences, Foster City, California, USA) and 1 µg RNA per 20 µl reaction.

#### Amplification

qPCR was performed with 9 ng cDNA, Taqman™ Gene Expression Master Mix (#4369016, ThermoFisherScientific, Waltham, Massachusetts, USA) and the respective TaqMan™ probes (Applied Biosystems, Foster City, California, USA). The following probes were used: Hypoxanthine Phosphoribosyltransferase 1 (HPRT1; Hs002800695_m1), CHI3L1 (Hs01072228_m1), GAP43 (Hs00967138_m1), APOE (Hs00171168_m1), TP53 (Hs01034249_m1) and CDKN1A (Hs00923894_m1). All reactions were carried out in a 96-well reaction plate (#N8010560, Applied Biosciences), covered with MicroAmp™ optical adhesion film (#4311971, Applied Biosciences, Foster City, California, USA) on a QuantStudio™ 3 Real Time PCR System (RRID:SCR_018712, ThermoFisherScientific, Waltham, Massachusetts, USA). ≥ 2 independent experiments with each having ≥ 2 technical replicates were performed. PCR reactions were checked by omission of templates and by melting curve and agarose gel electrophoresis. Standard curves were generated for each gene and the amplification was 85-115% efficient. Relative quantification of gene expression was determined by comparison of threshold values. All results were normalized to HPRT1 as the housekeeping gene.

### Antibody and recombinant blocking in vitro experiments

PDGCLs S24-TurboGFP, T269-TurboGFP and BG5-TurboGFP were cultured and singularized as described before. Cells were resuspended in PDGCL media as described before, however without growth factors but supplemented with glucose (#G7021-1KG, Sigma, part of Merck, Darmstadt, German) and cells seeded into a precoated uClear^®^ 96-well plate. Growth factor reduced Matrigel^®^ (#356231, Corning, Corning Inc., Corning, New York, USA) dissolved in PDGCL media was used for coating. Recombinant CHI3L1, CHI3L1 antibody or IgG1 Ctrl antibody were added immediately after cell seeding. Subsequently, cells were cultured under standard conditions (20% O2, 5% CO2, 37°C). All further procedures were described before. ≥ 9 images were acquired and analyzed per condition in each of ≥ 2 independent experiments. The Mann-Whitney U test was used.

### TP53 overexpression

#### Cloning

For functional analysis open reading frames (ORFs) of *TP53* wt and the mutation variants R175H (CGC>CAC), R248W (CGG>TGG) and R248Q (CGG>CAG) lacking a stopcodon were generated in a universal entry vector (pDONR221) for the use with the Gateway™ recombination system (Thermo Fisher Scientific, Waltham, Massachusetts, USA). After sequence validation, the ORFs were recombined in into a lentiviral expression vector rwpLX305-GW-Flag-CT-IRES-NeoR (Cellular Tools GPCF DKFZ, Heidelberg, Germany). The vector adds a short immunogenic Flag-Tag at the C-terminal end of the TP53 proteins to test for expression of the recombinant protein and couples a Neomycin resistance marker for selection of transduced and expressing cells via an IRES sequence.

#### Virus production and infection

For generation of lentiviral particles, HEK293FT cells (#R70007, Thermo Fisher Scientific, Waltham, Massachusetts, USA) were co-transfected with the lentiviral TP53 expression constructs and 2nd generation viral packaging plasmids VSV.G (kind gift from Tannishtha Reya, Addgene plasmid # 14888, RRID:Addgene_14888, http://n2t.net/addgene:14888) and psPAX2 (kind gift from Didier Trono, Addgene plasmid #12260, RRID:Addgene_12260, http://n2t.net/addgene:12260). 48h after transfection, virus containing supernatant was removed and cleared by centrifugation (5min/500g). The supernatant was passed through a 0.45 μm filter (#760517, Ahlstrom, Helsinki, Finland). U87 (#HTB-14, RRID:CVCL_0022, ATCC, Manassas, Virginia, USA) cells were transduced with lentiviral particles at 70% confluency in the presence of polybrene (TR-1003-G, Merck, Darmstadt, Germany). 24 h after transduction virus containing medium was replaced byG-418 sulfate (#M3118.0050, GENAXXON bioscience, Ulm, Germany) containing selection media.

### Target staining and TM quantification in patient tumor tissues

#### General preparation of slides

To validate the degree of connectivity and underlying markers we chose the three patients with high and three with low connectivity signature scores from which specimens were available at the Department of Neuropathology. Several consecutive sections were generated using the HM 355S automated microtom (#905200, ThermoFisherScientific, Waltham, Massachusetts, USA) and mounted on Superfrost slides (#J1800AMNZ, ThermoFisherScientific, Waltham, Massachusetts, USA). Subsequent drying was allowed for 30 min on a 37° C hot plate followed by baking for 10 min in a 75° C oven.

#### CHI3L1 and nestin staining

CHI3L1 and nestin expression was detected using the ultraView DAB protocol on the automated VENTANA^®^ BenchMark ULTRA platform (Roche, Basel, Switzerland).

After pretreatment involving deparaffinization CHI3L1 antigen retrieval CC1 solution (#05279801001, Roche, Basel, Switzerland) was applied for 32 min. Slides were subsequently incubated with anti-CHI3L1 antibody for 32 min. To detect nestin expression no heat induced epitope retrieval (HIER) was performed and slides were incubated with with anti-nestin antibody for 32 min. VENTANA^®^ standard signal amplification and ultra-wash was followed by counterstaining with Hematoxylin II (#790-2208, Roche, Basel, Switzerland) and blueing reagent (#760-2037, Roche, Basel, Switzerland) for 4 min each. Slides were removed from the staining platform, washed with tap water and rinsed with deionized water. After staining, all specimens were immersed in a series of ethanol (EtOH) solutions (#20821.330, VWR, part of Aventor, Radnor, Pennsylvania, USA) of increasing concentrations until 100% and Xylol (#534056-4L, Sigma, part of Merck, Darmstadt, Germany). Eukitt^®^ (#6.00.01.0001.06.01.01, ORSAtec GmbH, Bobingen, Germany) was used for mounting.

#### Hematoxylin-Eosin (HE) staining

Slides were prepared as described before. For dewaxing and rehydration sections were passed through xylol (#9713.3, Roth, Karlsruhe, Germany) and decreasing concentrations of EtOH (#200-678-6; Fisher Scientific, Waltham, Massachusetts, USA) until the solution evenly flowed across the slide. Staining with Mayeŕs hematoxylin solution consisting of 0.1% hematoxylin (#1.04302.0100, Merck, Darmstadt, Germany), 0.02% sodium iodate (#6525; Merck, Darmstadt, Germany), 5% potassium aluminum sulfate (#8896.1; Roth, Karlsruhe, Germany), 5% chloralhydrate (#K318.1; Roth, Karlsruhe, Germany) and 0.1% citric acid (#3958.1; Roth, Karlsruhe, Germany) for 1 min was followed by blueing in running tap water for 3 min. Slides were incubated in eosin solution consisting of 10% Eosin G (#7089.2, Roth, Karlsruhe, Germany) and 2 drops of glacial acetic acid (#3738.1; Roth, Karlsruhe, Germany) in 70% EtOH (#200-678-6, Fisher Scientific, Waltham, Massachusetts, USA) for 30 s and subsequently rinsed in aqua bidest. Mounting of HE sections was done as described before.

### Quantification of TMs in FFPE patient samples

For image analysis three 500 x 500 pixel regions in each patient sample were selected based on number of nuclei (100 ± 20), nestin positivity and adjacency to denser tumor tissue. Then TMs were measured manually in these regions using Fiji. There were 20-84 TMs measured per image with a total of n = 898.

#### Image analysis of patient tissue

All slides were scanned at 20x resolution using an Axioscan Z1 slide scanner (RRID:SCR_020927, Zeiss, Jena, Germany). Zen 2.6 Blue Edition ® software (RRID:SCR_013672, Zeiss, Jena, Germany) was used to globally adjust the copies of original photomicrographs for white and black balance. Photomicrographs were additionally cropped, rotated and resampled to allow alignment with other stainings.

#### Alignment of nestin, CHI3L1 and HE staining

The procedure for scanning CHI3L1 and HE stained consecutive sections was similar to Nestin stained sections. Zen 2.6 Blue Edition ® software (RRID:SCR_013672, Zeiss, Jena, Germany) was used to globally adjust the copies of original photomicrographs for white and black balance. Photomicrographs were additionally cropped, rotated and resampled to allow alignment with other stainings. Subsequent removal of background shadows at the tile edges of no-sample containing tiles was done using Zen 2.6 Blue Edition ® software (RRID:SCR_013672, Zeiss, Jena, Germany)

#### Histoscoring of CHI3L1

A histoscore was used to assess the quantity of the CHI3L1 staining intensities of both global tumor tissue level but also of 500 x 500 pixel CHI3L1 crops aligned with the nestin crops, which had been independently selected before by a blinded person. A histoscore was used to assess the quantity of the CHI3L1 staining intensities of both global tumor tissue level but also of 500 x 500 pixel CHI3L1 crops perfectly aligned with the nestin crops, which had been independently selected before by a blinded person.

Histoscoring is a widely used semiquantitative classification of the staining intensity of heterogeneously stained tissues. Technically, the staining intensity of each individual cell is assigned to a scaled rating: 0 (negative), 1 (low), 2 (moderate), and 3 (high). A weighted histoscore is calculated by the formula:

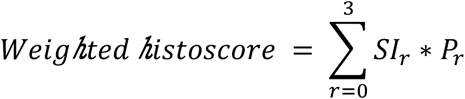

where *r* represents the rating of staining intensity; *SI_r_* represents the staining intensity of cell with *r*; *P_r_* represents the percentage of cells with *r* in the whole sample.

Based on this, the maximum score being reached is 300 (if 100% of cells have a high intensity) and the minimum score is 0 (if 100% of cells do not stain). All ratings were performed by a board-certified neuropathologist (Department of Neuropathology, University Hospital Heidelberg, Germany).

### Patient cohorts for validation of connectivity signature

#### TCGA (RRID:SCR_003193, https://www.cancer.gov/tcga, (33)) cohort

The RNA-Seq gene expression matrix, somatic mutation information, CNV information and clinical data of TCGA diffuse glioma samples were downloaded from UCSC Xena (RRID:SCR_018938, http://xena.ucsc.edu). We obtained 146 samples from TCGA GB cohort and 502 samples from TCGA lower grade glioma cohort. We further investigated *IDH* mutation status and chromosome 1p/19q co-deletion status in all samples. Finally, we obtained 230 *IDH* wt samples, 176 *IDH* mutant with 1p/19q co-deletion samples, 241 *IDH* mutant without 1p/19q co-deletion samples and one sample without clear classification. The 230 *IDH* wt samples were subjected for connectivity signature validation and survival analysis.

#### CGGA (RRID:SCR_018802, http://www.cgga.org.cn, (34)) cohort

We downloaded clinical data and RNA-Seq gene expression matrix of 325 GB samples from the CGGA webpage, of which 141 samples had *IDH* wt and intact 1p/19q status. These 141 samples were subjected for connectivity signature validation and survival analysis.

#### IvyGAP (RRID:SCR_005044, (35)) cohort

We obtained RNA-Seq gene expression matrix and corresponding laser micro-dissected anatomic structure information of 73 samples derived from 10 GB patients. Connectivity signature scores were calculated in each sample.

#### Gene Expression Profiling Interactive Analysis (RRID:SCR_018294, http://gepia.cancer-pku.cn, GEPIA, (36))

We downloaded the CHI3L1 gene expression level (transcripts per million [TPM]) of RNA-Seq data from GEPIA, which contains 31 tumor types from TCGA and related normal tissue samples from the genotype-tissue expression (GTEx).

### Molecular classification of TCGA RNA-seq

The TCGA and CGGA *IDH* wt GB samples were classified into three expression subtypes (i.e., mesenchymal, classical and proneural) by single sample GSEA analysis-based classification as described in (19) (ssGSEA, R codes from (19)). The fragments per kilo base per million mapped reads (FPKM) expression matrix was used as input for ssGSEA and 100,000 permutations was performed to obtain p values for each subtype. Each sample was assigned to the subtype with the smallest p value.

### Patient survival analyses

Connectivity signature scores were calculated in samples from TCGA and CGGA using AddModuleScore function in Seurat (RRID:SCR_007322, v.3.1.5). The TCGA/CGGA samples were assigned into three groups by Q1, Q2-Q3 and Q4 of connectivity scores. Kaplan-Meier survival analysis in the three groups using overall survival times, and Cox proportional hazards regression analysis with age, connectivity signature scores and overall survival times were performed with the survival (RRID:SCR_021137, v.3.1-12) and survminer (RRID:SCR_021094, v.0.4.2).

### Statistical analyses

A p value of p < 0.05 was generally considered significant. The p value of mean comparison between two groups was obtained by Mann-Whitney U test using ggpubr (RRID:SCR_021139, v.0.4.0). Pearson correlation coefficients were calculated using ggpubr (RRID:SCR_021139, v.0.4.0). Among the recurrent non-synonymous mutated genes in at least 5% TCGA GB patients (27 genes), the connectivity signature score related mutated genes were identified using wilcox.test function in R. Multiple testing was adjusted and obtained FDR using p.adjust function in R.

