## Supplement for "A connectivity signature for glioblastoma"

Ling Hai[1,2,3,4]*, Dirk C Hoffmann[2,3,4]*, Henriette Mandelbaum[2], Ruifan Xie[2], Jakob Ito[2], Erik Jung[2,3], Sophie Weil[2,3], Philipp Sievers[5,6], Varun Venkataramani[2,3,7], Daniel Dominguez Azorin[2], Kati Ernst[8,9], Denise Reibold[2], Rainer Will[10], Mario L. Suvà[11], Christel Herold-Mende[12], Felix Sahm[5,6], Frank Winkler[2,3], Matthias Schlesner[1,13], Wolfgang Wick[2,3], and Tobias Kessler[2,3#]

[1]Bioinformatics and Omics Data Analytics, German Cancer Research Center (DKFZ), Heidelberg, Germany
[2]Clinical Cooperation Unit Neurooncology, German Cancer Consortium (DKTK), German Cancer Research Center (DKFZ), Heidelberg, Germany
[3]Department of Neurology and Neurooncology Program, National Center for Tumor Diseases, Heidelberg University Hospital, Heidelberg, Germany
[4]Faculty of Biosciences, Heidelberg University, Heidelberg, Germany
[5]Department of Neuropathology, Institute of Pathology, University Hospital Heidelberg, Heidelberg, Germany
[6]Clinical Cooperation Unit Neuropathology, DKTK, DKFZ, Heidelberg, Germany
[7]Department of Neuroanatomy, Institute for Anatomy and Cell Biology, Heidelberg University, Heidelberg, Germany
[8]Hopp Children's Cancer Center at the NCT Heidelberg (KiTZ), Heidelberg, Germany
[9]Division of Pediatric Neurooncology, DKTK, DKFZ, Heidelberg, Germany
[10]Genomics and Proteomics Core Facility, DKTK, DKFZ, Heidelberg, Germany
[11]Broad Institute of Harvard and MIT, Cambridge, MA 02142, USA; Department of Pathology and Center for Cancer Research, Massachusetts General Hospital and Harvard Medical School, Boston, MA 02114, USA
[12]Department of Neurosurgery, Heidelberg University Hospital, Heidelberg, Germany
[13]Biomedical Informatics, Data Mining and Data Analytics, Faculty of Applied Computer Science and Medical Faculty, University of Augsburg, Augsburg, Germany

* Equally contributing first authors
### Corresponding author: Tobias Kessler MD, Neurology Clinic and Neurooncology Program at the National Center for Tumor Diseases & DKTK, DKFZ, Im Neuenheimer Feld 400, D-69120 Heidelberg, Germany. Phone: +49 6221 56 7075, Fax: +49 6221 56 7554,

#### Supplementary figures

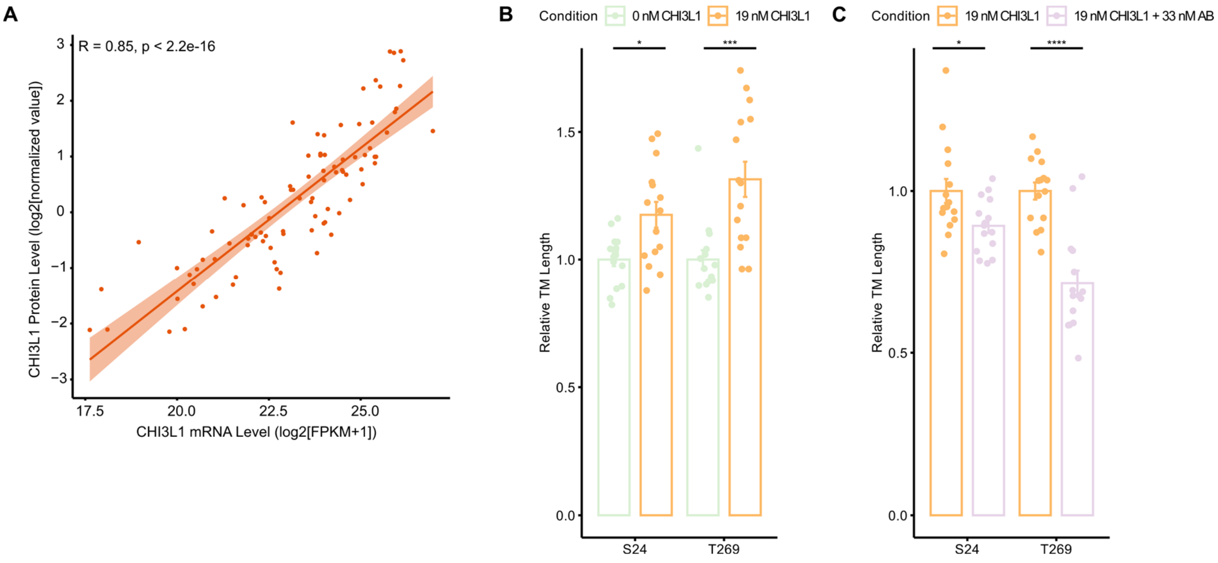


**Supplementary Figure 1:** Effects of recombinant CHI3L1 and antibody blocking on connectivity. **A,** Scatter plot showing correlation between CHI3L1 mRNA expression level (log2(FPKM+1)) and protein levels in 93 *IDH* wt GB samples from Wang et al. (1). Pearson correlation test was used to calculate correlation coefficients and p values. **B.** Bar plot of relative TM length per live cell of crops after administration of recombinant CHI3L1 protein compared to control in two PDGCLs. Two replicates per PDGCL. **C.** Bar plot of the relative TM lengths per live cell of crops after administration of recombinant CHI3L1 compared to recombinant CHI3L1 together with a CHI3L1 blocking antibody in two PDGCLs. A dot represents a crop. P value was calculated by Wilcoxon test.

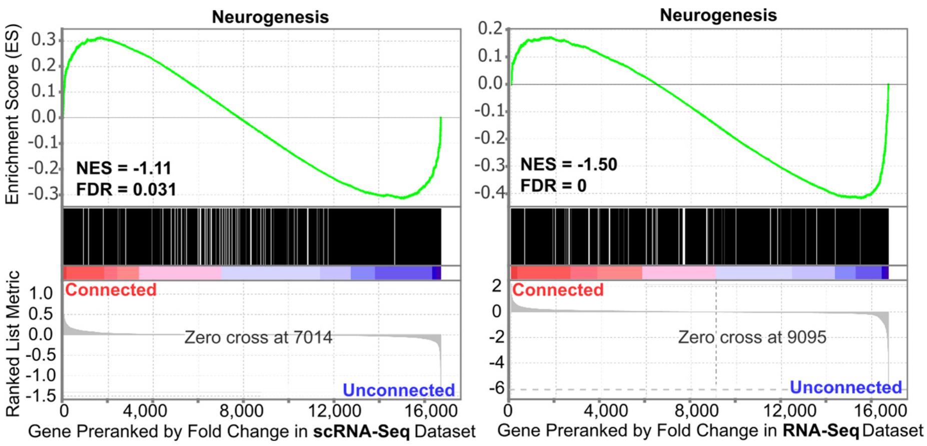


**Supplementary Figure 2.** Gene set enrichment analysis in 16,759 genes pre-ranked by fold changes between SR101$^{high}$ and SR101$^{low}$ samples for the neurogenesis gene set. Left, scRNA-Seq data from PDGCL xenografted mouse models. Right, RNA-Seq data from PDGCL xenografted mouse models. NES: normalized enrichment score, FDR: false discovery rate.

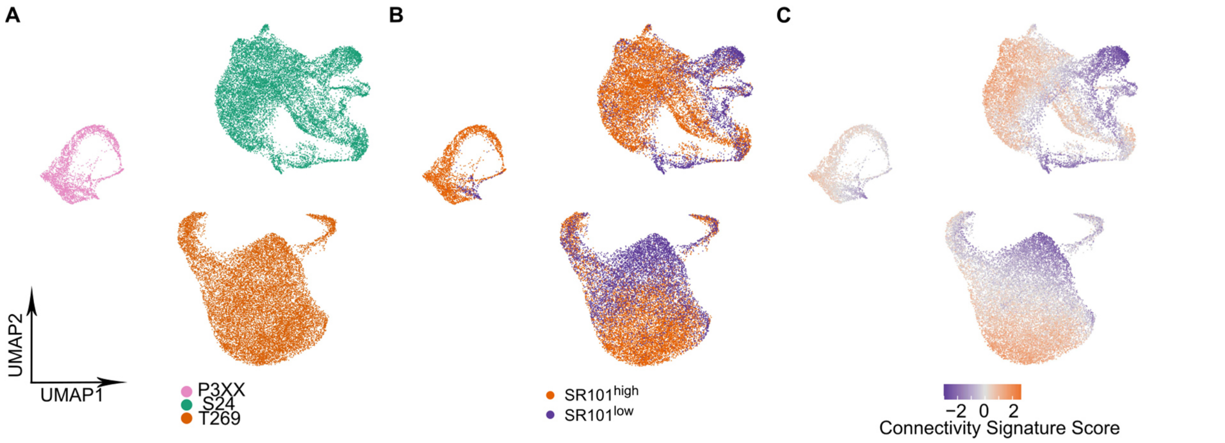


**Supplementary Figure 3.** UMAPs in scRNA-Seq of PDGCL xenograft models and oxygen condition experiment in PDGCLs. **A-C,** UMAPs of cells in three PDGCL xenograft mouse models without anchoring integration. **A,** colored by the derived PDGCL. **B,** colored by SR101 sorting. **C,** colored by connectivity signature scores. Scores were scaled, centered and winsorized at -3 and 3 across cells.

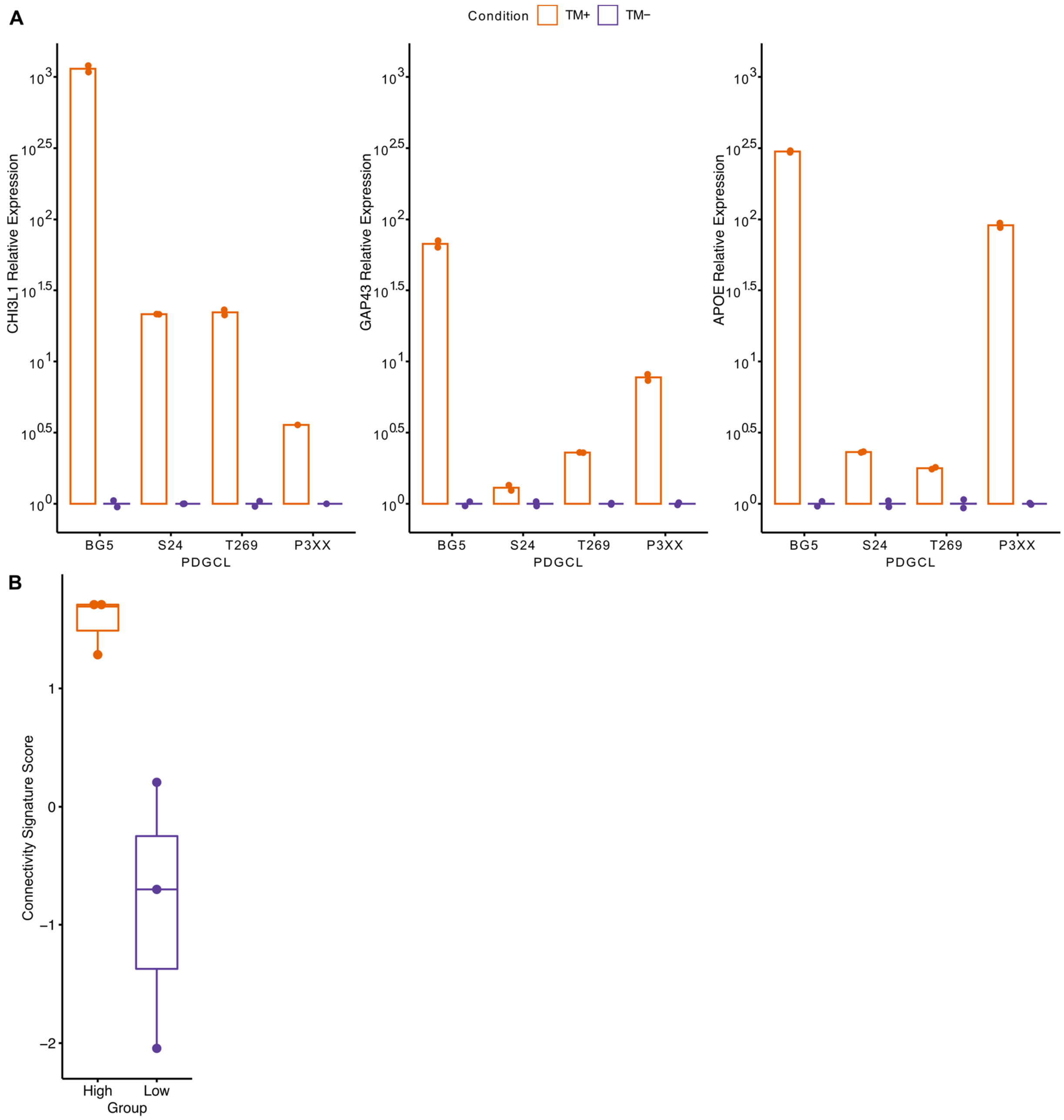


**Supplementary Figure 4.** Connectivity in cultivated cell lines. **A.** Box plot of relative gene expression measured by qPCR of *CHI3L1*, *GAP43* and *APOE* under TM+ condition compared to TM- condition in four PDGCLs. **B.** Box plot of connectivity signature scores in six patients.

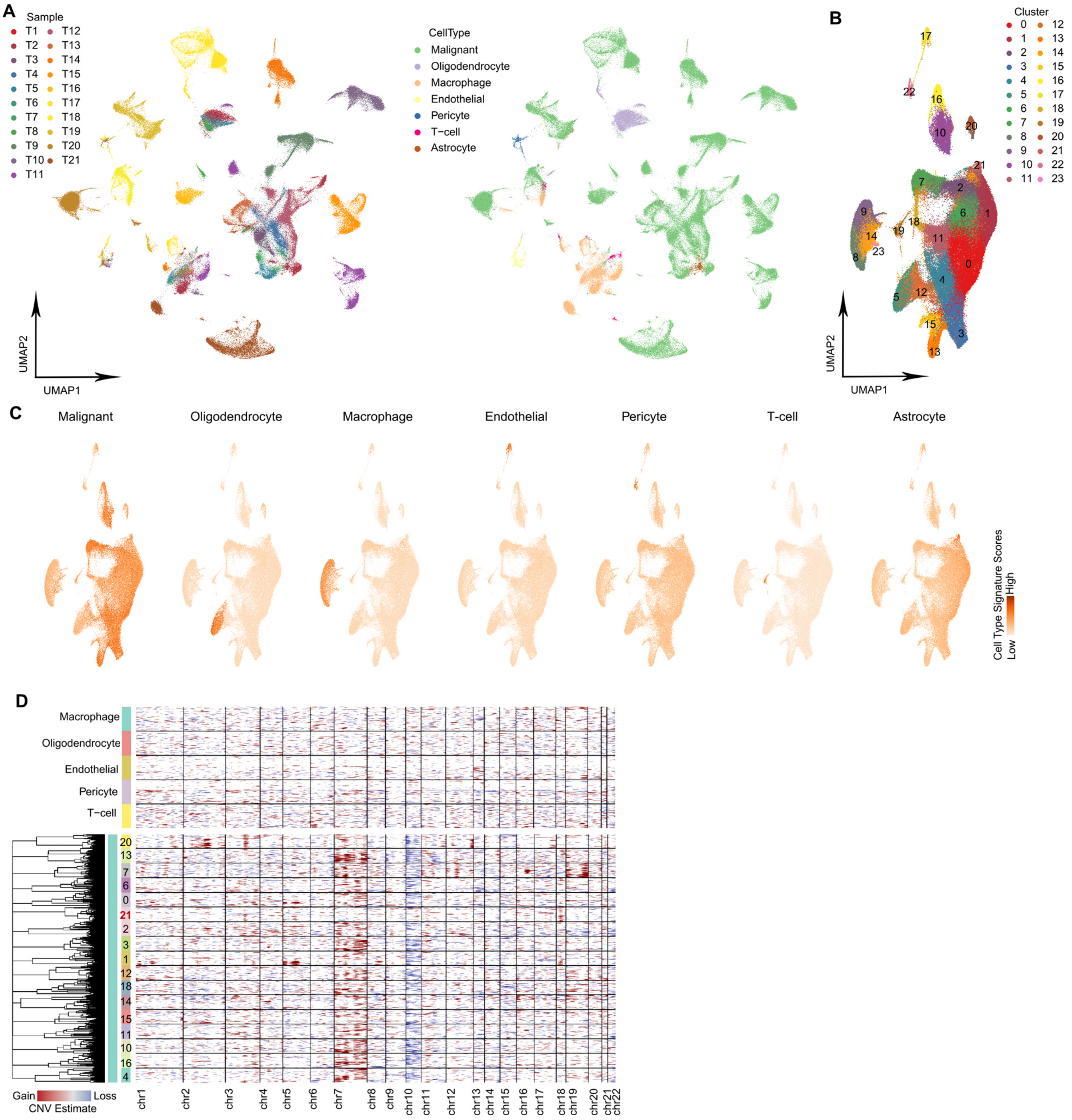


**Supplementary Figure 5.** snRNA-seq of 21 GBM patient samples. **A,** UMAPs of cells from 21 patient samples without anchoring integration. Left, colored by patient. Right, colored by malignant and non-malignant cell types. **B,** UMAP of patient samples with anchoring integration. Cells were colored by unsupervised clusters. 24 clusters were obtained. **C,** UMAPs of cell type signature scores of malignant and each non-malignant cell types separately. **D,** CNVs in non-malignant cell types and malignant clusters. Cluster 21 marked in red indicates the non-malignant astrocyte cluster.

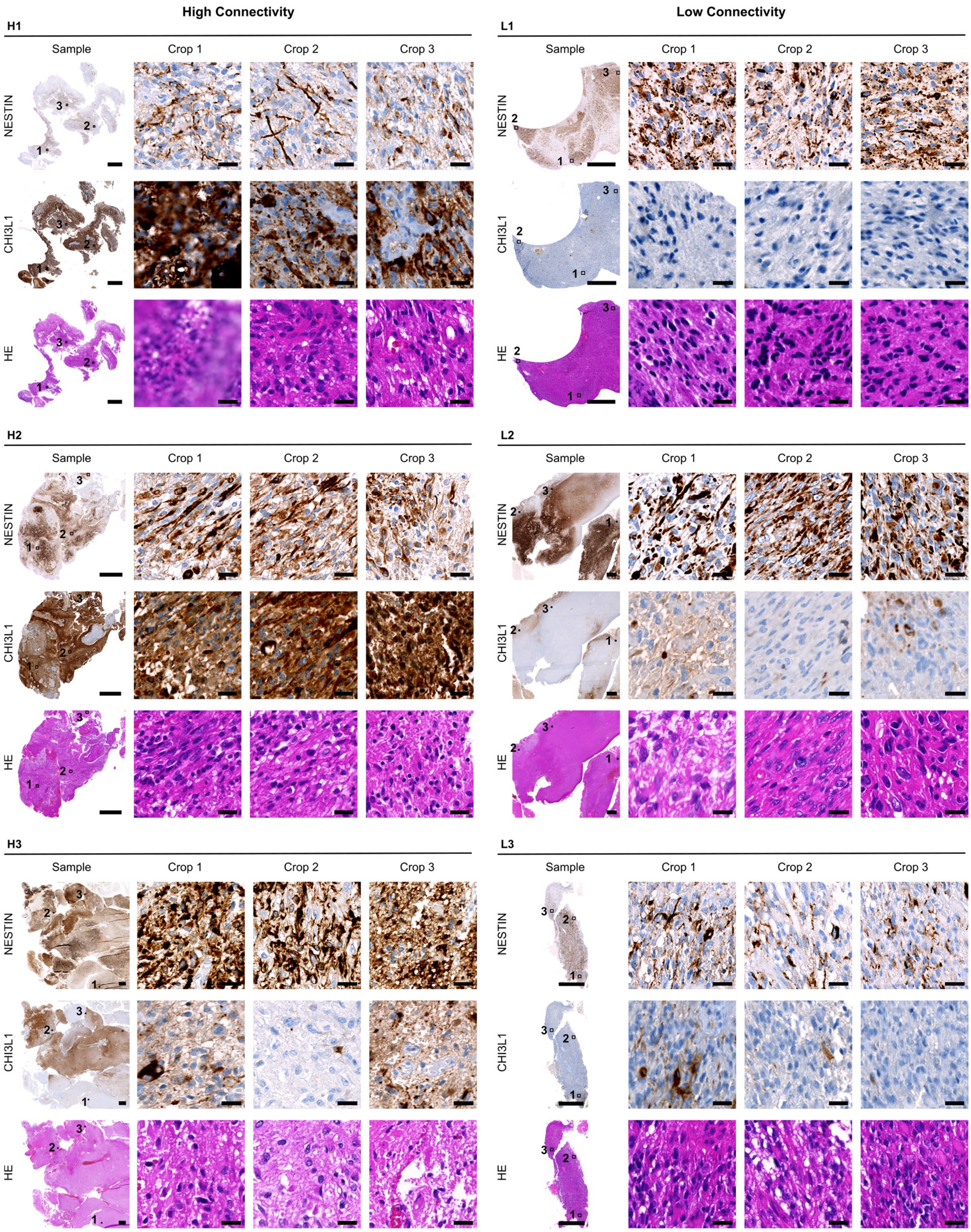


**Supplementary Figure 6.** Nestin, CHI3L1, and HE staining images of six GBM patients. Left column shows three patients with high connectivity signature scores (H1, H2 and H3), right column shows three patients with low connectivity signature scores

(L1, L2 and L3). In each patient, row shows staining of nestin, CHI3L1 or HE; column shows image with different size and regions. In the “Sample” column, three marked regions are the crops that zoomed in in the next three columns (“Crop1”, “Crop2” and “Crop3”), scale bar is 1000 μm. The scale bar of crops is 20 μm.

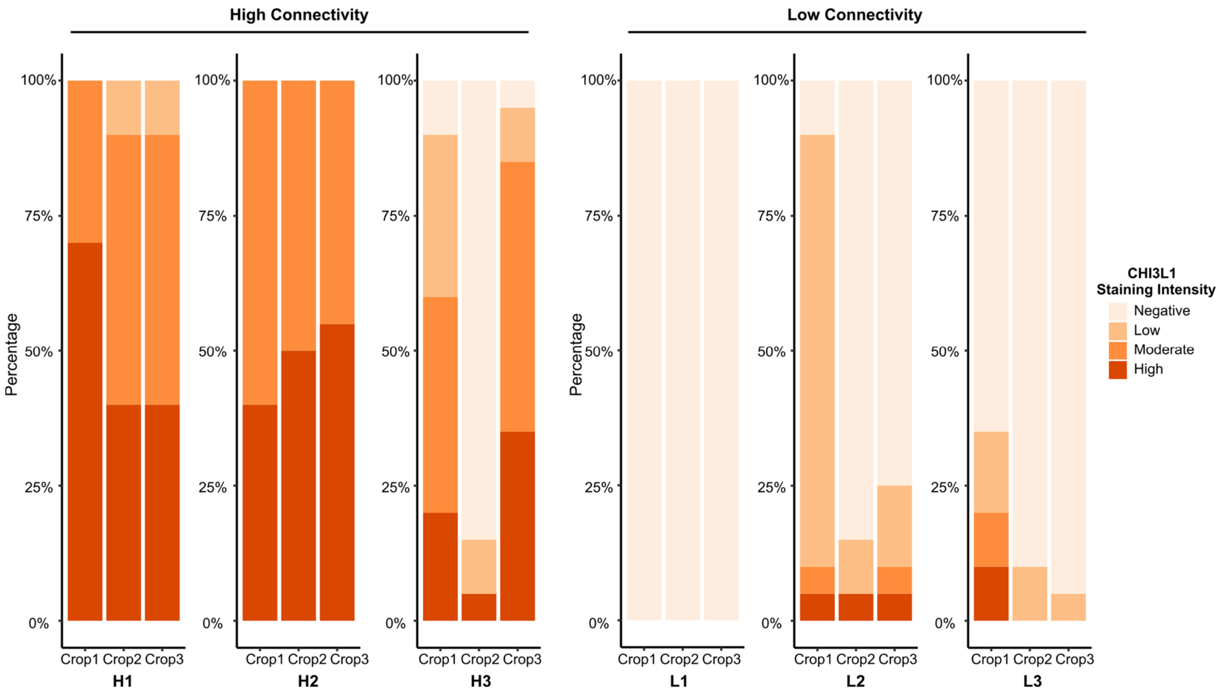


**Supplementary Figure 7.** The frequency of CHI3L1 staining intensity of cells in image crops shown in **Supplementary Figure 6**.

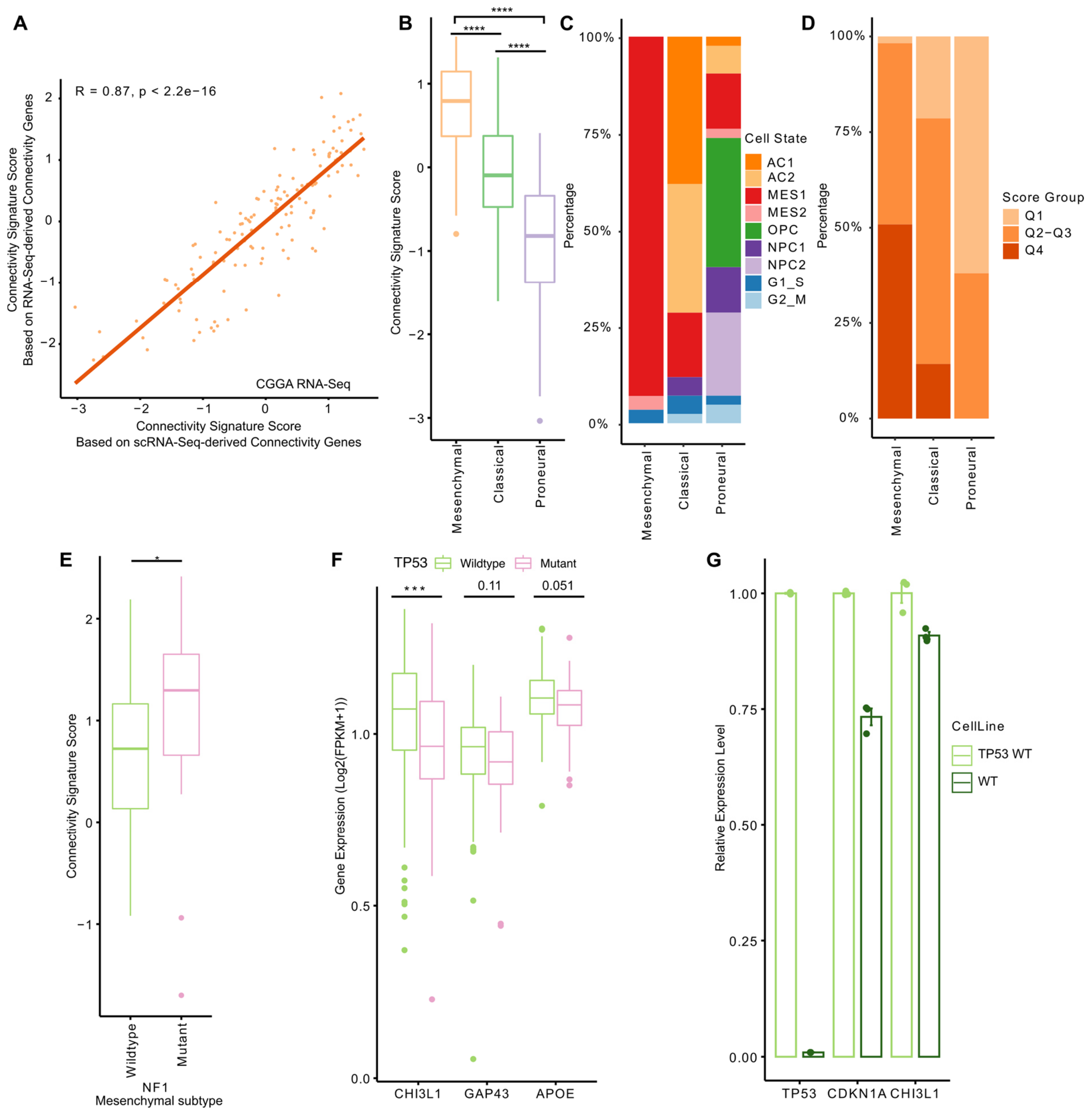


**Supplementary Figure 8.** Connectivity signature scores in TCGA and CGGA cohorts. **A,** Scatter plot showing connectivity signature scores based on connectivity genes derived from scRNA-Seq and RNA-Seq datasets in 141 CGGA *IDH* wt GB samples. Correlation coefficients and p values were calculated by Pearson correlation test. **B,** Boxplot of connectivity signature scores of three expression subtypes (57 mesenchymal, 42 classical and 42 proneural) in CGGA samples. **C,** Frequency of dominant cell states in each expression subtype. **D,** Frequency of connectivity signature score groups in each expression subtype. Connectivity signature scores grouped by first and last quartiles into Q1, Q2-Q3 and Q4. **E,** Box plot of connectivity signature scores in NF1 wt (n = 62) and NF1 mutant patients (n = 17) from TCGA IDH wt GBM mesenchymal subtype. **F,** Box plot of *CHI3L1*, *GAP43* and *APOE* gene expression (Log2(FPKM+1)) in TP53 wt (n = 173) and TP53 mutant patients (n = 57) from the TCGA IDH wt GBM cohort. **G,** Bar plot of relative gene expression of *TP53*, *CDKN1A* and *CHI3L1* in the U87 TP53 wt (WT) against U87 with overexpressing TP53 wt (TP53 WT) by qPCR. **D-G,** 141 CGGA glioblastoma samples. **H-J,** 417 TCGA IDH mutant glioma samples. **A, B, E,** connectivity scores were scaled, centered and winsorized at -3 and 3 across samples. **E, F, B,** P values were calculated by Wilcoxon test. *, $p < 0.05$; **, $p < 0.01$; ***, $p < 0.001$.

**A**

Cox Proportional Hazards Regression Survival Analysis

| | Multivariate analysis with Age, Gender and Subtype | |
|---|---|---|
| *n* = 230 | Exp. coef. (95% int.) | *p*-value |
| Connectivity Signature Score | 1.303 (1.058-1.604) | 0.0128 |

**B**

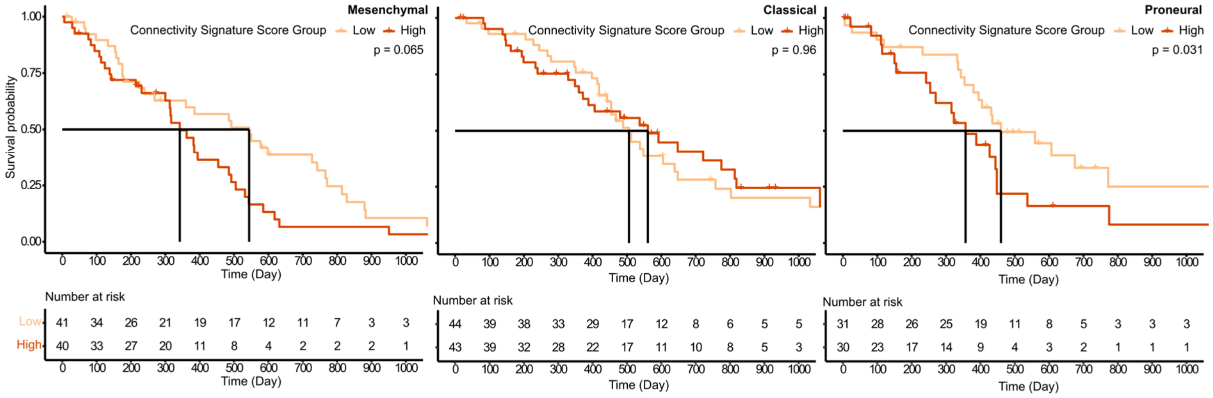


**C**

Cox Proportional Hazards Regression Survival Analysis

| | Multivariate analysis with Age, Gender and Subtype | |
|---|---|---|
| *n* = 230 | Exp. coef. (95% int.) | *p*-value |
| CHI3L1 Expression Level | 2.867 (0.986-8.334) | 0.053 |

**D**

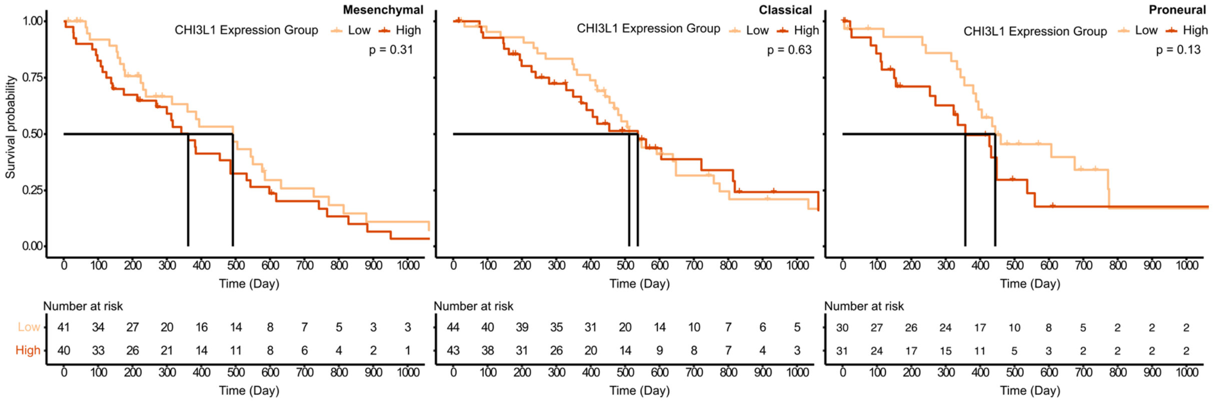


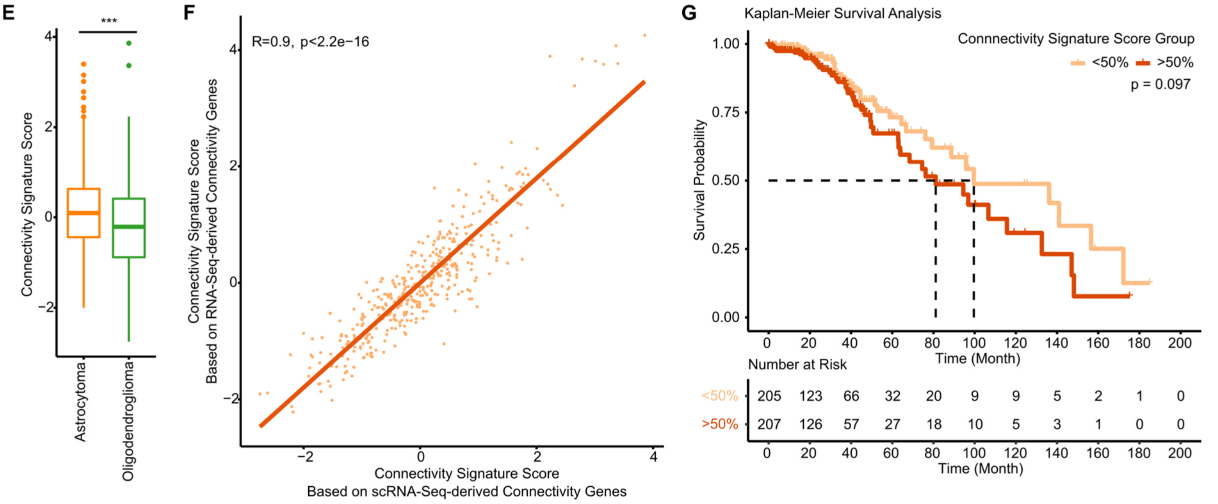


**Supplementary Figure 9.** Survival analysis of connectivity signature score in TCGA cohort. **A**, Cox proportional hazards regression survival analysis with connectivity signature scores, adjusting for ages, genders, and expression subtypes, in 230 TCGA IDH wt GB samples. **B,** Kaplan-Meier survival analysis with two connectivity signature score groups (dividing by median of connectivity signature scores) in each expression subtype separately. **C**, Cox proportional hazards regression survival analysis with *CHI3L1* expression levels (log2[FPKM+1]), adjusting for ages, genders, and expression subtypes, in 230 TCGA IDH wt GB samples. **D,** Kaplan-Meier survival analysis with two *CHI3L1* expression groups (separating by median of connectivity signature scores) in each expression subtype separately. **E,** Box plot of connectivity signature scores in 241 astrocytoma and 176 oligodendroglioma from TCGA *IDH* mutant samples. **F,** Scatter plot showing connectivity signature scores based on connectivity genes derived from scRNA-Seq and RNA-Seq datasets. Correlation coefficients and p values were calculated by Pearson correlation test. **G,** Kaplan-Meier survival analysis of TCGA *IDH* mutant samples according to two connectivity signature score groups divided by median.

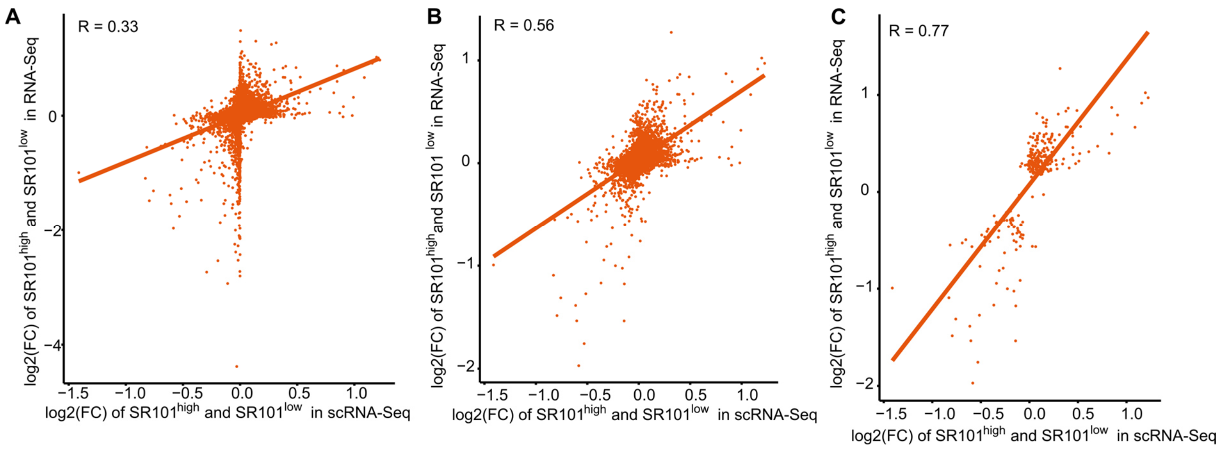


**Supplementary Figure 10.** Correlation of the fold changes (FC) of $SR101^{high}$ and $SR101^{low}$ samples in scRNA-Seq and RNA-Seq datasets of PDGCL xenografted mouse models. **A,** plotting with 16,759 genes. **B,** plotting with the genes which were expressed in more than 10% cells of scRNA-Seq dataset (6,984 genes). **C,** plotting with the genes which were significantly regulated between $SR101^{high}$ and $SR101^{low}$ samples in both datasets (303 genes, adjust p-value < 0.05).

#### Supplementary tables

**Supplementary Table 1.** Properties of PDGCLs.

| Name | IDH | MGMT | Methylation classifier | scRNA-Seq of xenografted mouse model | RNA-Seq of xenografted mouse model | scRNA-Seq of *in vitro* model |
|---|---|---|---|---|---|---|
| **S24** | WT | M | RTK I | + | + | + |
| **T269** | WT | M | RTK II | + | + | + |
| **P3XX** | WT | M | RTK II | + | - | + |
| **BG5** | WT | M | RTK I | - | - | + |

WT, wildtype; M, promoter methylated; U, promoter unmethylated; RTK, receptor tyrosine kinase; +, used in the experiment; -, not use in the experiment.

**Supplementary Table 2.** Detailed description of 40 connectivity genes upregulated in highly connected cells.

| GENE | CORRELATION TO Glioma/Connectivity/Tumor microtubes (TMs) |
|---|---|
| **AGT/ Angiotensinogen** | • **AGT** und **CHI3L1** are part of a 14-gene-signature that defines a high-risk group of GB patients and is associated with immune response and mesenchymal subtype (2)<br>• **Angiotensin II** promotes axonal elongation of postnatal rat retinal explants, dorsal root ganglia neurons in vitro and axonal regeneration of retinal ganglion cells after optic nerve crush in vivo (3)<br>• Treatment of nondifferentiated NG108-15 (neuroblastoma x glioma) cells with Angiotensin II induces morphological differentiation of neuronal cells and outgrowth of neurites. This is correlated with an increased level of polymerized tubulin and microtubule-associated protein MAP2c. (4)<br>• **AGT** mRNA and protein are expressed by nontumoral astrocytes and GB cells (5)<br>• data suggests that GB cells have the potential to produce **AGT**/prorenin/renin and **Angiotensin** I (5)<br>• Serum proteomic analysis using iTRAQ method: When comparing GB tumors close (SVZ+) and distantly (SVZ-) located to the subventricular zone (SVZ), AGT (Angiotensinogen) and CHI3L1 are significantly altered tissue proteins in GBM tumors close to the subventricular zone. The SVZ is a major site of neurogenesis (6) |
| **ANXA2/ Annexin A2** | • Knockdown of **AnnexinA2** decreases migration capacity of U87MG and U373MG glioma cells in vitro (7)<br>• Knockdown of **AnnexinA2** reduces glioma growth and invasion in vivo and reduces proliferation and angiogenesis in glioma tissue (8)<br>• **ANXA2** is elevated in harvested pseudopodia of U87 glioma cells (9) |
| **APOE** | • **APOE** is significantly upregulated in serum of patients with GB tumors close to the subventricular zone (SVZ+) indicating its potential role in tumor proliferation in GB (6)<br>• **APOE** is a major component of LDL and supports cell proliferation and anti-apoptotic pathways (6)<br>• **APOE** positive cells have a higher index in the calculated connectivity score and seem to be strongly involved in TM formation and maintenance (10) |
| **ATF3** | • **ATF3** is induced in sensory and motoneurons in the spinal cord following peripheral nerve injury (11)<br>• **ATF3** as a transcription factor (TF) and GAP43 as a gene are components of the neuron intrinsic regeneration-associated gene response (RAG) that occurs after nerval injury (12) |

| | |
|---|---|
| | • the RAG program is activated during injury in the peripheral nervous system (PNS) but remains weak in the central nervous system (CNS). By applying a peripheral lesion as a "conditioning lesion", the RAG program is induced, and central axons can regenerate (12)<br>• **ATF3** is a regeneration associated TF that regulates the RAG and promotes neurite outgrowth and axon regeneration (12)<br>• **ATF3** promotes neurite outgrowth (13)<br>• both **ATF3** and GAP43 overexpression in dorsal root neurons (DRGs) lead to a higher number of neurons that extend long processes compared to control (13)<br>• **ATF3** transgenic mice were constructed with a comparable expression level of **ATF3** as in peripheral nerve injured DRGs (= elevated). The regeneration capacity is stronger in **ATF3** transgenic mice. Moreover, the amount of GAP43 positive fibers distal from the nerve injury doubled in **ATF3** transgenic mice compared to controls. However, GAP43 gene expression is not affected in **ATF3** transgenic mice.(14)<br>• **ATF3** knockdown inhibits growth of glioma xenograft tumors in vivo (15) |
| **ATP1B2** | • high **ATP1B2** expression is associated with poor prognosis in GBM (16)<br>• **ATP1B2** knockdown inhibits GB cell proliferation and colony formation in vitro (16)<br>• **ATP1B2** expression in clearly associated with GB growth in vivo (16) |
| **B2M** | None found |
| **C1orf61** | None found |
| **CALM1/ Calmodulin 1** | • **Calmodulin** is involved in regulation of cell proliferation (17)<br>• **Calmodulin** is involved in the cytoskeleton of the cell and associates with multiple microtubule-associated proteins (17)<br>• **Calmodulin** is a calcium binding protein that promotes GBM invasion by supporting invadopodium formation (18)<br>• **Calmodulin** is associated with poor prognosis and aggressiveness in GB (18) |
| **CD63 and TIMP1** | • **TIMP-1** mRNA levels correlate with higher tumor grade and lower TIMP-1 expression levels predict longer survival of patients with GB (19)<br>• **TIMP-1** and **CD63** are expressed in close molecular proximity in GB tissue and may interact with one another (20)<br>• Co-expression of **TIMP-4** and **CD63** is an independent prognostic factor for poor survival in GB patients (21) |
| **CEBPD** | • **CEBPD** - together with CEBPB and STAT3- is a component of master transcriptional regulators of the |

| | |
|---|---|
| | mesenchymal transition and strongly associated with the extent of necrosis and poor prognosis in GB (22) |
| **CHI3L1/ YKL-40** | • **CHI3L1** shows significant upregulation in mesenchymal-like cells compared to proneural cells (2)<br>• **CHI3L1** shRNA knockdown reduces Matrigel invasion in U87MG cells (23)<br>• **CHI3L1** inhibition leads to reduction of MMP2 expression in western blotting and enzyme activity (23). MMPs facilitate TM formation throughout the brain. (24)<br>• Adhesion ability of U87MG cells was significantly reduced by CHI3L1 suppression (23)<br>• **CHI3L1** sh-transfected U87MG cells show an increase of stress fibers (23)<br>• **CHI3L1** overexpression increases invasion and adhesion capacity in U373MG cells (23)<br>• Overexpression of **CHI3L1** in U373MG cells causes reorganization and morphological changes of the F-actin cytoskeleton (23)<br>• Vinculin is a component of mature focal adhesions and staining with anti-vinculin in control cells stayed only in the center of the cell whereas staining in **CHI3L1** overexpressing U373MG cells spreads to both the leading edge and the center (23)<br>• **CHI3L1** overexpression is associated with increased ability of colony forming in soft agar<br>and is associated with drug resistance (23)<br>• **YKL-40 (CHI3L1)** is associated with the mesenchymal subtype with high glioma grade, chemotherapy **resistance** and glioma aggressiveness (25)<br>• **CHI3L1 and CLU** seem to be a clear marker for a mesenchymal cell type in GB that is also described in astrocytic like markers (26)<br>• **CHI3L1** is associated with loss of chromosome 10 and poor survival (27)<br>• **CHI3L1** is one of the most overexpressed genes in GB and is a patient serum marker that correlates with glioma grade (28)<br>• **YKL-40** regulates VEGF expression and both trigger angiogenesis in GB in vitro (29)<br>• **YKL-40** plays a role in the proliferation of connective tissue cells and connective tissue turnover in chondrocytes and synoviocytes (30) |
| **CITED1** | None found |
| **CLU** | • **CLU** is overexpressed in GB (31) |
| **CNN3/ Calponin3 / acidic Calponin** | • **Calponins** are actin-filament stabilizing molecules (32)<br>• **Calponin 3/acidic Calponin** is expressed in the brain, regulates dendritic spine morphology and density presumably through regulation of the actin cytoskeleton (33) |

| | |
|---|---|
| | • acidic- calponin- induced spines can establish functional glutamatergic synapses (33) |
| **CRYAB/ Crystallin Alpha B** | • CRYAB is strongly expressed by highly migratory GB cells (34) |
| **CST3** | None found |
| **CYR61/ CCN1** | • among patients with totally resected tumors, **CCN1** expression correlates negatively with PFS and OS (35)<br>• **CCN1** interacts with S1P and regulates glioma invasion and extracellular matrix adhesion thereby contributing to GBM invasiveness (36) |
| **DNAJB1** | None found |
| **FABP7** | • **FABP7** is involved in proliferation and invasion of GB cells (37)<br>• **FABP7** is vital to lipid metabolism in particularly infiltrative slow cycling cells (SCC) that support invasion, chemoresistance and tumor proliferation in GB (38)<br>• Inhibition of **FABP7** in SCCs reduces their stress survival capacity and leads to overall increased survival (38)<br>• Higher **FABP7** expression correlates with poorer patient survival (38) |
| **FEZ1** | • **FEZ1** associates with microtubules with its COOH-terminal region in PC12 cells (39)<br>• **FEZ1** interacts with cytoskeletal filaments within signal cascades important for growth cone and neurite outgrowth (39)<br>• agnoprotein colocalizes with microtubules in infected JCV cells and induces the dissociation of **FEZ1** from microtubules (39)<br>• **FEZ1** plays a critical role in neuronal development and acts together with Kinesin1 to transport proteins important for neurite outgrowth and synaptic formation/function (40)<br>• proteins that are specifically enriched in Kinesin1 and **FEZ1** immunoisolated samples (vesicles from the rat brain cytosol fraction analyzed by mass spectrometry) are classified by Ingenuity Pathway Analysis as crucial for: neurotransmission, synaptic transmission, potentiation of synapses and branching of neurons (40)<br>• cargos of Kinesin 1 are within others: GluR2/GRIA2 and commonly identified in the shared proteins that are upregulated in immunostained **FEZ1** cells (40)<br>• phosphorylation of **FEZ1** is mainly done by microtubule-affinity-regulating -kinases (MARKS/ PAR-1) (40). The phosphorylation by MARKS influences its task in presynaptic protein trafficking (40)<br>• UNC-64 is a known cargo of **FEZ1** and is transported through axons of ventral nerve cord neurons in wild type worms. When worms are par1-deficient/mutated, axonal |

| | |
|---|---|
| | transport of UNC-64/UNC76 is damaged and abnormal axonal aggregates of both proteins and cell body retentions are visible (40)<br>• SNB1 is a marker for presynaptic specializations. Par1 mutant worms show significantly lower SNB1 densities compared to the wt worms. Moreover, larger interpunctata distances between (GFP-SNB1 puncta) - that are representing presynaptic organization defects - are present in par1 mutant worms (40)<br>• **FEZ1** interacts with the c-Jun-N-terminal-kinase-binding-protein (JIP1) (41)<br>• **FEZ1** and JIP1 bind to microtubule-based motor kinesin 1 and lead to kinesin motor activity and activation of the JNK signaling pathway (41)<br>• the JNK signaling pathway (and its receptor Grindelwald) is required for TM formation (24)<br>• JNK signaling leads to the post transcriptional upregulation and accumulation of matrix metalloproteinases MMPs (24)<br>• MMPs contribute to a successful infiltration of TMs throughout the brain (24) |
| **GADD45B** | • **GADD45B** and members of the Wnt pathway family genes are upregulated in response to neuronal ischemic damage and seem to play a neuroprotective role (42) |
| **GAP43** | • **GAP43** is a proven main driver of TM formation and maintenance (43)<br>• knockdown of **GAP43** reduces intracellular calcium waves (ICWs) and the proportion of astrocytes extending multiple TMs *in vivo* (43)<br>• **GAP43** deficiency leads to decreased tumor size in MRI and better survival (43) |
| **GAPDH** | • **GAPDHs** regulation as a response to hypoxia is not an absolute phenomenon and **GAPDH** is a valid alternative to a housekeeping gene in human Gliomas (44) |
| **GFAP** | • **GFAP** serum levels correlate with tumor volume in high grade gliomas (45) (46)<br>• member of cytoskeletal protein family and astroglial marker (47) |
| **HOPX** | • **HOPX** is a transcription factor and marker of astrocytic lineage in the dorsal subventricular zone (48)<br>• **HOPX** seems to be a regulator of self-renewal capacity in glioma cancer stem cells that impacts neurogenesis (49) |
| **ID3** | • expression of **ID1-3** in astrocytic brain tumors correlates with its grade of malignancy (47)<br>• **ID3** might play a role in EGFR-ID3 mediated induction of angiogenic cytokines in GB tumor cells (50) |
| **MT2A/ Metallothionein-II/MT-II** | • Bt2-cAMP can overcome MAG (myelin associated glycoprotein)- induced inhibition of axonal regeneration after nerval injury (51) |

| | |
|---|---|
| | • genes involved in the cAMP pathway that enables axonal regeneration are (51) metallothioneins (MT I and II)<br>• **MT II** is the major isoform of MT I that is found in the CNS (51)<br>• **MT2A** promotes regeneration of the optic nerve in vivo after injury (51)<br>• a statistically significant increase of axonal regeneration in the optic nerve is detected in MTII injected animals compared to controls injected with saline when quantified by staining for GAP43 positive axons at 500µm (51)<br>• axonal density was also significantly higher for MTI/II injected animals (51) |
| **NDUFB2** | • **NDFUB2** is part of the GABAergic synapse SuperPath in Path Cards *https://pathcards.genecards.org/card/gabaergic_synapse*<br>• **NDFUB2** is part of the endocannabinoid signaling Pathway together with GRIA2+GRIA3 |
| **NEK6** | • **NEK6** has been described of being overexpressed in several cancer types (52)<br>• **NEK6** increases **STAT3** phosphorylation at SER727 (52)<br>• **STAT3** is an oncogenic transcription factor and its phosphorylation plays an important role in oncogenesis and proliferation of GB and is shown to correlate with tumor grade (53) |
| **NMB** | • **NMB** is member of a four gene panel correlating with poor survival of GB patients (54)<br>• **NMB** stimulates the clonal growth of C6 GB rat cells and increases cytosolic Calcium levels in single C6 cells (55) |
| **PTN/ Pleiotrophin** | • **PTN** plays a role in the guidance of neuroblast migration and neurite outgrowth in neurodevelopment (56)<br>• **Pleiotrophin** is -together with integrins and neurotrophins - involved in the formation of cellular protrusions of NPCs and TMs in glioma (56) |
| **S100A16** | • **S100A16** is a calcium binding protein and is astrocyte specific (57)<br>• **S100A16** protein is expressed in human GB U373 cells and translocates from the nucleolus to the cytoplasm after calcium stimulation (57) |
| **SOCS3** | • **SOCS3** is highly expressed in primary GB tumor samples and SOCS3 positive cells localize near necrosis and blood vessels within the tumor tissue (58)<br>• **SOCS3** regulates self-renewal in GB stem cells and correlates with poor survival in GB patients (59) |
| **SPARC and SPARCL1** | • **SPARC** is secreted by astrocytes. SPARC interacts with beta3integrin and controls levels of synaptic AMPARs (60)<br>• **SPARC** drives the ability of developing synapses to regulate plasticity during synapse maturation: **SPARC-KO** mice show elevated AMPAR levels and greater AMPAR mediated currents (60) |

| | |
|---|---|
| | • neurons of WT mice were treated with TTX (tetrodotoxin) to induce activity blockade in synapses. This was compensated by neurons by increasing AMPAR (GluR1) levels in order to increase synaptic strength. Neurons in **SPARC** depleted mice were not able to increase their AMPAR levels and remained at a same level than before TTX treatment.<br>When "rescued" with recombinant **SPARC** protein, these neurons were able to increase their GluR1 levels. This means neurons cultured during **SPARC** absence maintain their potential to increase AMPAR levels when **SPARC** is added.<br>**SPARC** keeps the synapses within a useful "working range". This enables them to modulate postsynaptic developmental modifications according to activity (60)<br>• **SPARC** and Hevin are secreted by astrocytes and work together in synaptogenesis<br>(61, 62)<br>• **SPARC** inhibits specifically hevin induced synaptogenesis and is a synapse regulatory protein (61, 62)<br>• **SPARC** negatively and Hevin positively regulate synapse formation and morphology (61, 62)<br>• **SPARC** upregulates the expression of membrane type 1-matrix metalloproteinase (MT1-MMP) and matrix metalloproteinase-2 (MMP-2) transcripts (63)<br>• MMP1 and MMP2 protein accumulation on glial cells are dependent on TM formation (and Fz1 mediated Wg signaling) (24)<br>• **SPARCL1** positively correlates with tumor grade in glioma (64)<br>• **SPARCL1** protein is also expressed in NPCs of the subventricular zone and a binding partner of above described pleiotrophin (56) and both drive glioma invasion by Rho/ROCK signaling pathway activation (65) |
| **TAGLN2** | • **TAGLN2** positively correlates with tumor grade, worse prognosis, and the mesenchymal subtype in glioma (66)<br>• Gene silencing of **TAGLN2** in U87MG and U251 glioma cells decreased tumor growth in vitro and in vivo (66)<br>• **TAGLN2** is an actin-binding protein and a key regulator in cytoskeleton remodeling and modulation of cellular processes and overexpressed in various cancer types (67) |
| **TMSB10** | None found |
| **TNFRSF12A/ FN14** | • visual stimulation induces a unique gene program in excitatory neurons of the dorsal lateral geniculate nucleus of the thalamus in mice and **FN14** is the most inducible molecule specific to the excitatory population (68)<br>• **FN14** regulates pre- and postsynaptic morphology in the dorsal lateral geniculate nucleus of the thalamus during visual stimulation (68) |

| | |
|---|---|
| | • **Fibroblast growth factor-inducible 14 (FN14)** is relatively higher expressed in migrating glioma cells compared to stationary cells and levels are elevated in GB tissue compared to normal brain tissue (69)<br>• TWEAK (tumor necrosis factor-like weak inducer of apoptosis) is the only known receptor for **FN14** and treatment of **FN14** positive cells with TWEAK stimulates migration, invasion, and resistance against chemo in GBM (69)<br>• **FN14** levels are elevated in recurrent GB (70)<br>• GBM PDX cells with TMZ resistance have higher **FN14** levels than their parental counterpart (70)<br>• investigations concerning the expression level of **FN14,** and clinical consequences showed that:<br>patients with higher **FN14** levels had shorter OS and DFS than those with lower expression (71)<br>• cell line data in the same study suggests that **FN14** might lead to resistance against TMZ and might predict sensitivity to TMZ (71) |
| **TUBB2A** | • Protein network analysis and differential gene expression analysis using GEO public database revealed TUBB2A as one of the top 5 most important genes within GB interacting networks (72)<br>• Microtubules, key participants in processes such as mitosis and intracellular transport, are composed of heterodimers of alpha- and beta-tubulins. The protein encoded by this gene is a beta-tubulin. |

**Supplementary Table 3.** The prediction performances of connectivity signature scores against SR101-sorted labels.

| **Metrics** | **245 RNA-Seq-derived connectivity genes** | **71 scRNA-Seq-derived connectivity genes** | **245 randomly generated genes** | **71 randomly generated genes** |
|---|---|---|---|---|
| Accuracy | 0.79 | 0.83 | 0.49 | 0.49 |
| Sensitivity | 0.77 | 0.95 | 0.47 | 0.48 |
| Specificity | 0.83 | 0.58 | 0.53 | 0.51 |
| PPV | 0.90 | 0.82 | 0.67 | 0.67 |
| NPV | 0.64 | 0.84 | 0.32 | 0.32 |

PPV, positive predictive value; NPV, negative predictive value.

**Supplementary Table 4.** Clinical characteristics and quality measures of scRNA-Seq patients.

| ID | Age | IDH | MGMT | GBM meth. class | Primary/ recurrent | Median gene (n) | Median count (n) | Cell (n) | Malignant cell (n) |
|---|---|---|---|---|---|---|---|---|---|
| T1 | 69 | WT | U | RTK I | primary | 561 | 685 | 932 | 800 |
| T2 | 61 | WT | U | Mesenchymal | recurrent | 786 | 1116 | 16721 | 6423 |
| T3 | 68 | WT | U | RTK I | primary | 634 | 794 | 6634 | 6184 |
| T4 | 61 | WT | U | RTK II | primary | 943 | 1273 | 9029 | 7455 |
| T5 | 77 | WT | U | RTK II | primary | 767 | 1006 | 6175 | 3232 |
| T6 | 73 | WT | U | Mesenchymal | primary | 1020 | 1440 | 2744 | 1916 |
| T7 | 56 | WT | U | RTK II | primary | 643 | 846 | 5009 | 2757 |
| T8 | 80 | WT | U | Mesenchymal | primary | 584 | 679 | 1626 | 1552 |
| T9 | 67 | WT | U | Mesenchymal | primary | 1393.5 | 2071 | 15092 | 13737 |
| T10 | 64 | WT | U | Mesenchymal | primary | 1097 | 1653.5 | 11192 | 10573 |
| T11 | 44 | WT | M | N/A | primary | 1198 | 1762 | 14588 | 10918 |
| T12 | 66 | WT | M | RTK I | primary | 996 | 1381 | 15057 | 13751 |
| T13 | 54 | WT | U | RTK I | primary | 1231 | 1946 | 5165 | 4526 |
| T14 | 69 | WT | U | RTK II | primary | 1289 | 2053 | 11927 | 10795 |
| T15 | 53 | WT | M | RTK II | primary | 998 | 1310 | 11533 | 11081 |
| T16 | 43 | WT | U | RTK II | primary | 608 | 728 | 5830 | 5600 |
| T17 | 64 | WT | U | Mesenchymal | primary | 1347.5 | 1917 | 19668 | 16840 |
| T18 | 56 | WT | U | Mesenchymal | primary | 810 | 1102 | 8221 | 5140 |
| T19 | 55 | WT | U | RTK I | primary | 1300.5 | 1890 | 13450 | 10963 |
| T20 | 32 | WT | U | Mesenchymal | primary | 856 | 1055 | 15707 | 15140 |
| T21 | 55 | WT | U | RTK II | primary | 1912 | 3024.5 | 17144 | 13088 |

N, number; WT, wildtype; M, promoter methylated; RTK, Receptor tyrosine kinase; U, promoter unmethylated; censor 1, progressive disease; censor 0, stable disease
